# Preclinical Efficacy of IMM-BCP-01, a Highly Active Patient-Derived Anti-SARS-CoV-2 Antibody Cocktail

**DOI:** 10.1101/2021.10.18.464900

**Authors:** Pavel A. Nikitin, Jillian M. DiMuzio, John P. Dowling, Nirja B. Patel, Jamie L. Bingaman-Steele, Baron C. Heimbach, Noeleya Henriquez, Chris Nicolescu, Antonio Polley, Eden L. Sikorski, Raymond J. Howanski, Mitchell Nath, Halley Shukla, Suzanne M. Scheaffer, James P. Finn, Li-Fang Liang, Todd Smith, Nadia Storm, Lindsay G. A. McKay, Rebecca I. Johnson, Lauren E. Malsick, Anna N. Honko, Anthony Griffiths, Michael S. Diamond, Purnanand Sarma, Dennis H. Geising, Michael J. Morin, Matthew K. Robinson

**Affiliations:** Immunome, Inc., Exton, PA, U.S.A.; Department of Microbiology, Boston University School of Medicine and National Emerging Infectious Diseases Laboratories, Boston, MA, USA; Departments of Medicine, Molecular Microbiology, Pathology & Immunology, Washington University School of Medicine, St. Louis, MO 63110, USA

## Abstract

Using an unbiased interrogation of the memory B cell repertoire of convalescent COVID-19 patients, we identified human antibodies that demonstrated robust antiviral activity *in vitro* and efficacy *in vivo* against all tested SARS-CoV-2 variants. Here, we describe the pre-clinical characterization of an antibody cocktail, IMM-BCP-01, that consists of three unique, patient-derived recombinant neutralizing antibodies directed at non-overlapping surfaces on the SARS-CoV-2 spike protein. Two antibodies, IMM20184 and IMM20190 directly block spike binding to the ACE2 receptor. Binding of the third antibody, IMM20253, to its unique epitope on the outer surface of RBD, alters the conformation of the spike trimer, promoting release of spike monomers. These antibodies decreased SARS-CoV-2 infection in the lungs of Syrian golden hamsters, and efficacy *in vivo* efficacy was associated with broad antiviral neutralizing activity against multiple SARS-CoV-2 variants and robust antiviral effector function response, including phagocytosis, ADCC, and complement pathway activation. Our pre-clinical data demonstrate that the three antibody cocktail IMM-BCP-01 shows promising potential for preventing or treating SARS-CoV-2 infection in susceptible individuals.

**One sentence summary:** IMM-BCP-01 cocktail triggers Spike Trimer dissociation, neutralizes all tested variants *in vitro*, activates a robust effector response and dose-dependently inhibits virus *in vivo*.

## INTRODUCTION

With over 472 million cases and more than 6.1 million deaths worldwide (<u>Johns Hopkins</u> <u>University Coronavirus Resource Center)</u>, the SARS-CoV-2 pandemic continues to pose extraordinary health and economic challenges. The scientific community has mitigated this threat through the discovery and launch of a myriad of vaccines and therapeutics to prevent or treat infections. While the initial data for the spike (S) protein-directed vaccines have been impressive, the current rate of protection against variants of concern is decreasing, which was predicted to occur due to viral escape and patient immunodeficiency or immunosuppression (*1–6*).

As such, the discovery and development of effective antibody therapies for passive immunization with broad range of reactivity is likely to be an important alternative approach to vaccination. The use of convalescent plasma against SARS-CoV-2 initially yielded mixed results (*7, 8*), however a recent retrospective cohort study showed reduced mortality in treated patients (*9*) outlining the need for a more robust and more standardized antiviral antibody cocktail. A phase 3 clinical trial with Lilly’s Bamlanivimab was halted on the basis of data showing no improvement in clinical outcomes. An early Regeneron trial with a 2-antibody mixture was also paused based on a potential safety signal and an unfavorable risk/benefit profile (*10*). Nonetheless, subsequent data demonstrated that S-protein-directed antibodies can have significant efficacy and safety, and both the Lilly and Regeneron antibody cocktail candidates received Emergency Use Authorization (EUA) from the US FDA in November 2020, although the EUA for Bamlanivimab was later withdrawn and distribution of the Bamlanivimab/Etesevimab cocktail is now limited to areas where resistant variant frequency is below 5%. Another S-protein specific antibody, Sotrovimab, co-developed by Vir and GlaxoSmithKline, received an EUA in May 2021. As publicly reported, Vir is currently developing a second-generation antibody aimed for use as a combination with Sotrovimab. The study published by Regeneron demonstrated that both Regeneron 2-Ab cocktail and Vir’s VIR-7831 antibody generated escape mutants after seven and two passages *in vitro*, outlining a need for multiple neutralizing antibodies in a cocktail (*11*). Further, the recent SARS-CoV-2 variants of concern (VOC), Omicron (BA.1, BA.1.1, and BA.2), was shown to escape Regeneron and Lilly’s antibody cocktails, that led to the FDA’s decision to limit the use of bamlanivimab and etesevimab cocktail and REGEN-COV (casirivimab and imdevimab cocktail) to patients infected with susceptible variants (that are currently not detected in the US) (*12, 13*). The emergence of Omicron variants recently led the FDA to revise the EUA issued for another combination Evusheld (consists of tixagevimab and cilgavimab), and increase the dose due to loss of potency to BA.1 and BA.1.1(*14*). Finally, a new Lilly’s antibody bebtelovimab, that received an EUA in February of 2022, was demonstrated to retain activity against Omicron(*15*). However, earlier findings from monoantibody therapies confirm the need for a cocktail treatment to avoid generation of escape mutants. Therefore, an antibody cocktail with broad reactivity and limited possibility of escape to current and prospective VOC that consists of several antibodies to block the generation of escape mutants is an urgent, yet unmet, medical need.

Small molecule inhibitors (SMI), which target viral proteins other than S protein, are alternatives to antibody-based therapies that might not be affected by the current VOC. However, SMI have additional limitations, such as the requirement to inhibit patient’s CYP3A for a viral protease inhibitor PF-07321332 or a low enough dose to avoid the host DNA mutagenesis for a ribonucleoside analog molnupiravir(*16*). In addition, SMIs are associated with toxicity concerns that could limit clinical usefulness for some patient populations. (*17, 18*). Thus, the collateral effects and resistance patterns of SMI may need to be considered prior to patient dosing.

We previously reported the identification of a library of patient-derived antiviral antibodies (*19*). Based on published reports (*20*), we hypothesized that interrogation of such patient responses would identify rare immunoglobulins against epitopes that have a synergistic antiviral effect when combined and are resistant to mutational drift. In this report, we describe the pre-clinical efficacy of IMM-BCP-01, that we are planning to move to clinical trials. IMM-BCP-01 consists of three patient-derived antibodies, each selected for its own intrinsic antiviral neutralization and functional effector response activities against current isolates and prospective variants. These antibodies bind to three non-overlapping epitopes on the receptor binding domain (RBD) and, when combined, potently neutralize *in vitro* all tested viral variants, including Alpha, Beta, Gamma, Epsilon, Kappa, Delta, Mu, Omicron, and suppress viral spread in the lungs of infected animals *in vivo.* We observed a drop in viral load in the lungs of animals treated with our antibody cocktail in a dose-dependent manner. Each antibody demonstrates unique binding properties: IMM20190 has a composite epitope involving the ACE2 receptor binding ridge and an area adjacent to the receptor binding loop, IMM20184 binds avidly to two S proteins within the same trimer, and IMM20253 binds to a conserved epitope on the outer surface of the RBD. Our experiments show that three antibody cocktail IMM-BCP-01 has potent and broad antiviral activity in animals, which makes it a promising candidate for development for humans to combat infection with emerging SARS-CoV-2 variants.

## RESULTS

### Three antibody cocktail IMM-BCP-01 binds to conserved non-overlapping epitopes of S protein trimer leading to its re-organization and dissociation into S protein monomers

Using an unbiased interrogation of a previously described library of patient-derived antiviral antibodies (*19*), we identified three monoclonal antibodies (mAbs), IMM20190, IMM20184 and IMM20253, that had robust additive and synergistic combinatorial antiviral effects. Structural (**Fig 1**) and functional (**Fig 2-4**) studies of these antibodies revealed a unique mechanism of action of the IMM-BCP-01 cocktail (**Fig 5**).

**Figure 1.**
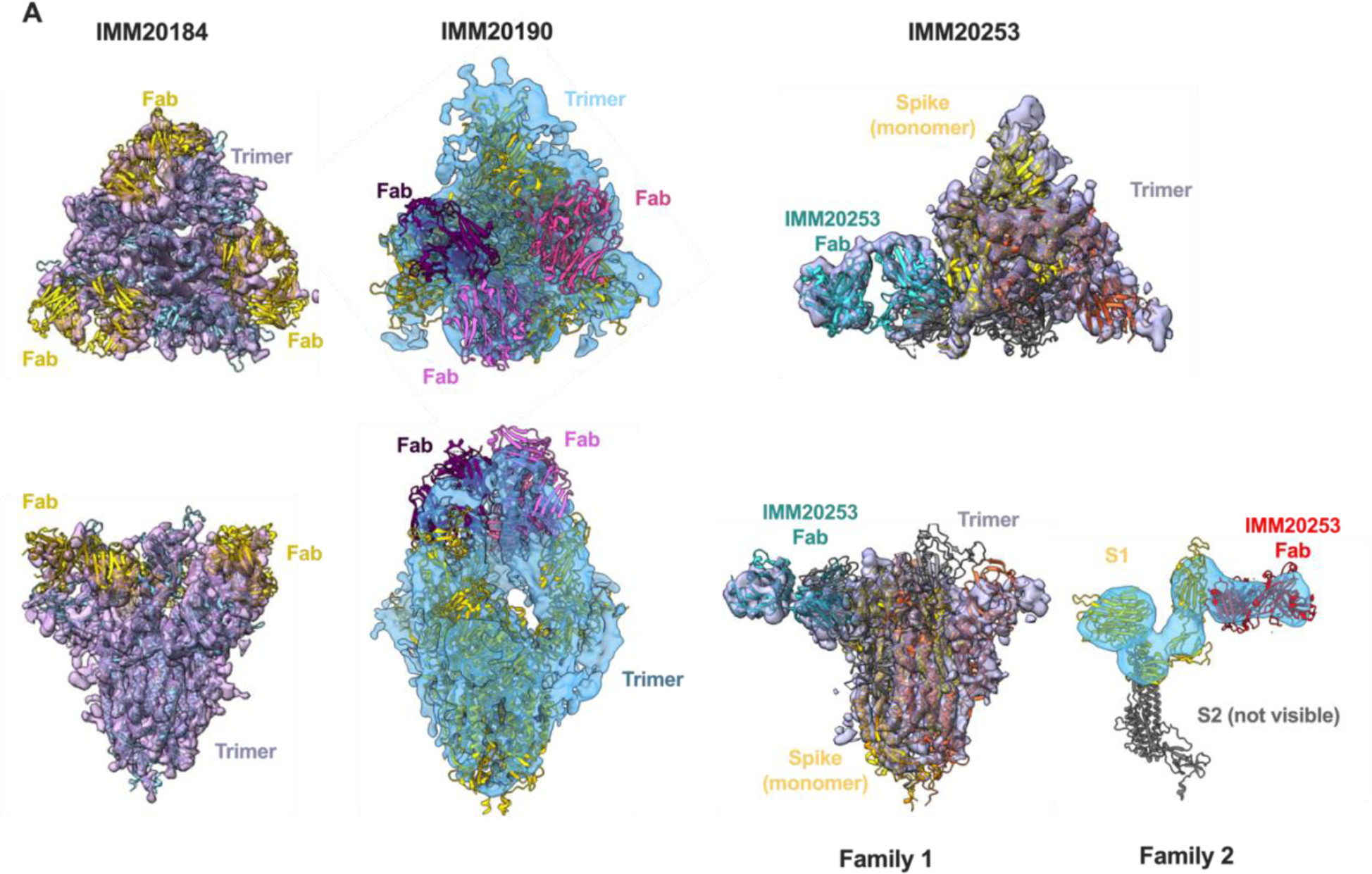
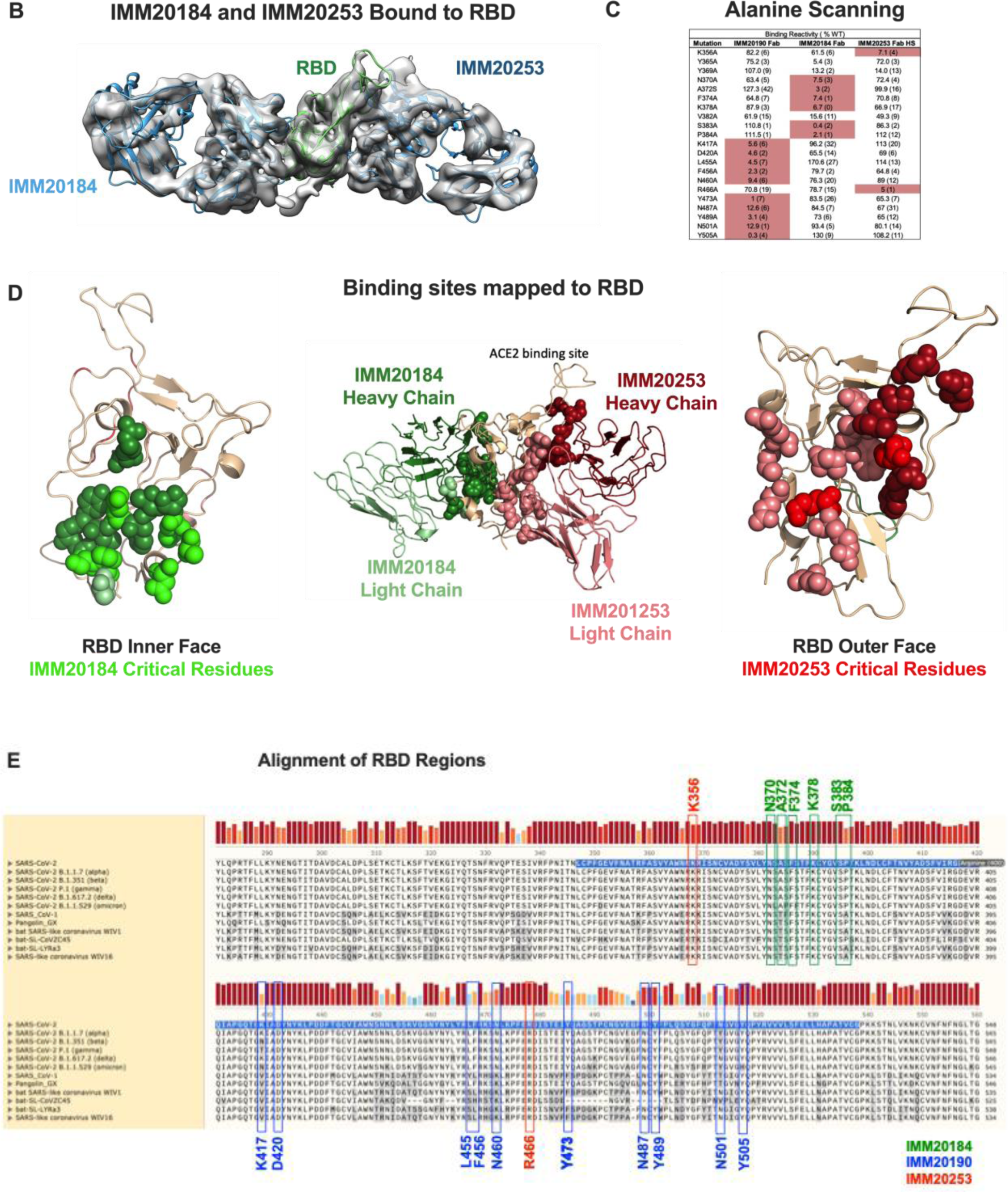
IMM20184, IMM190 and IMM20253 antibodies bind to conserved epitopes and disrupt Spike Trimer. (A) 3D reconstruction of Cryo-EM images of Spike Trimer complex with either IMM20184 (left), IMM20190 (middle) or IMM20253 (right) Fabs at ∼7Å resolution. Top-to bottom view (top) and side view (bottom) are shown. IMM20253 Fab binding to Trimer results in two families, Family 1 and 2. Models PDB:7E8C, PDB:6XLU, PDB:6XM5 or PDB:7NOH were fit to density in Chimera for the Spike Trimer and PDB:6TCQ for the Fabs. (B) 3D reconstruction of Cryo-EM images of a complex of RBD with simultaneously bound Fabs of IMM20184 and IMM20253 at ∼3.9Å. Fab model PDB:1M71 was fit to density in Chimera. (C) Critical residues of antibody epitopes identified as binding pattern to a library of single-point RBD mutants expressed on the cell surface. (D) Epitopes of IMM20184 and IMM20253 antibodies mapped on the RBD model PDB: 7A97. Epitopes of IMM20184 (green) and IMM20253 (red) antibodies and critical residues (bright green and red) are shown. (E) Alignment of Spike protein sequences from current and prior CDC VOCs, SARS-CoV-1 and closely related coronaviruses. Critical residues of IMM20190, IMM20184 and IMM20253 epitopes are shown in blue, green and red. Highlighted sequence indicates RBD.

**Figure 2.**
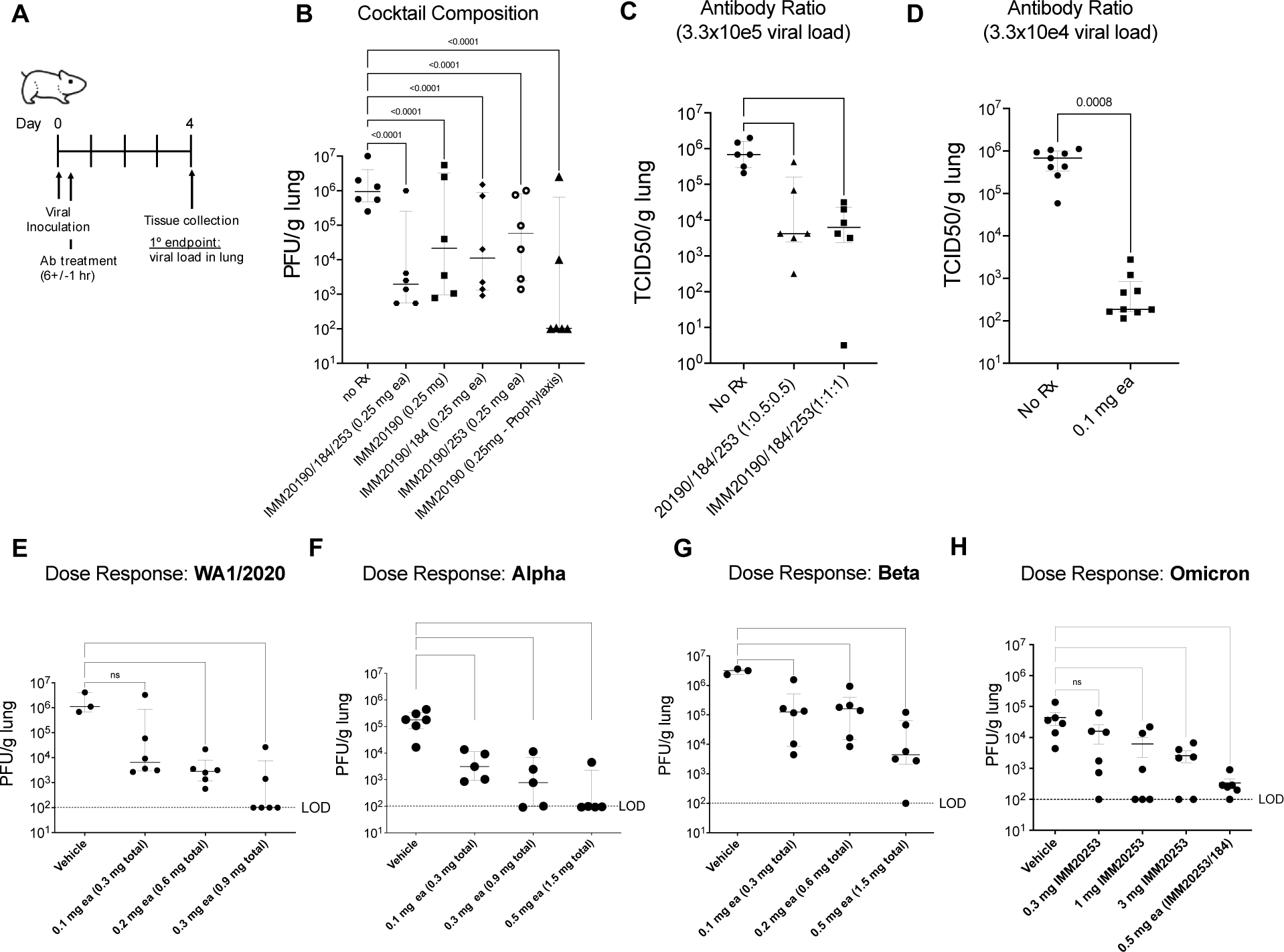
Three antibody combinations IMM20190/184/253 inhibits replication of non-adapted SARS-CoV-2 in lungs of infected animals. (A) All studies were carried out in Syrian Golden hamsters challenged with an intra-nasal inoculation of SARS-CoV-2 and treated with antibodies post-inoculation. Lungs were harvested at Day 4 and viral titers determined by either plaque forming or TCID50 assays. (B) Animals were infected with WA_CDC-WA1/2020 isolate and treated with single, double, or triple antibody cocktails, as noted, 6 hours post-inoculation. (C) Hamsters were challenged with 3.3 x 10^5^ TCID50 viral inoculation and treated with 3-Ab cocktail, at two different antibody ratios (1:1:1 or 1:0.5:0.5). (D) Hamsters were challenged with 3.3 x 10^4^ TCID50 viral inoculation and treated with 3-Ab cocktail at 1:1:1 ratio, at 0.1 mg dose each (0.3 mg total). (E -H) Hamsters were challenged with 10e4 PFU of WA1/2020 (E), Alpha (B.1.1.7) (F) Beta (B.1.351) (G), or Omicron (BA.1) (H) SARS-CoV-2 isolates after pre-treatment (Day -1) with different doses of 3-Ab cocktail IMM20190/184/253 at 1:1:1 ratio. Bar denotes median values. Error bars denote interquartile range. Statistical analysis in panel B is Two-Way ANOVA and in panels C-H is One-way ANOVA using Dunnet’s multiple comparisons test comparing to untreated group (No Rx) available in Prizm 9.

**Figure 3.**
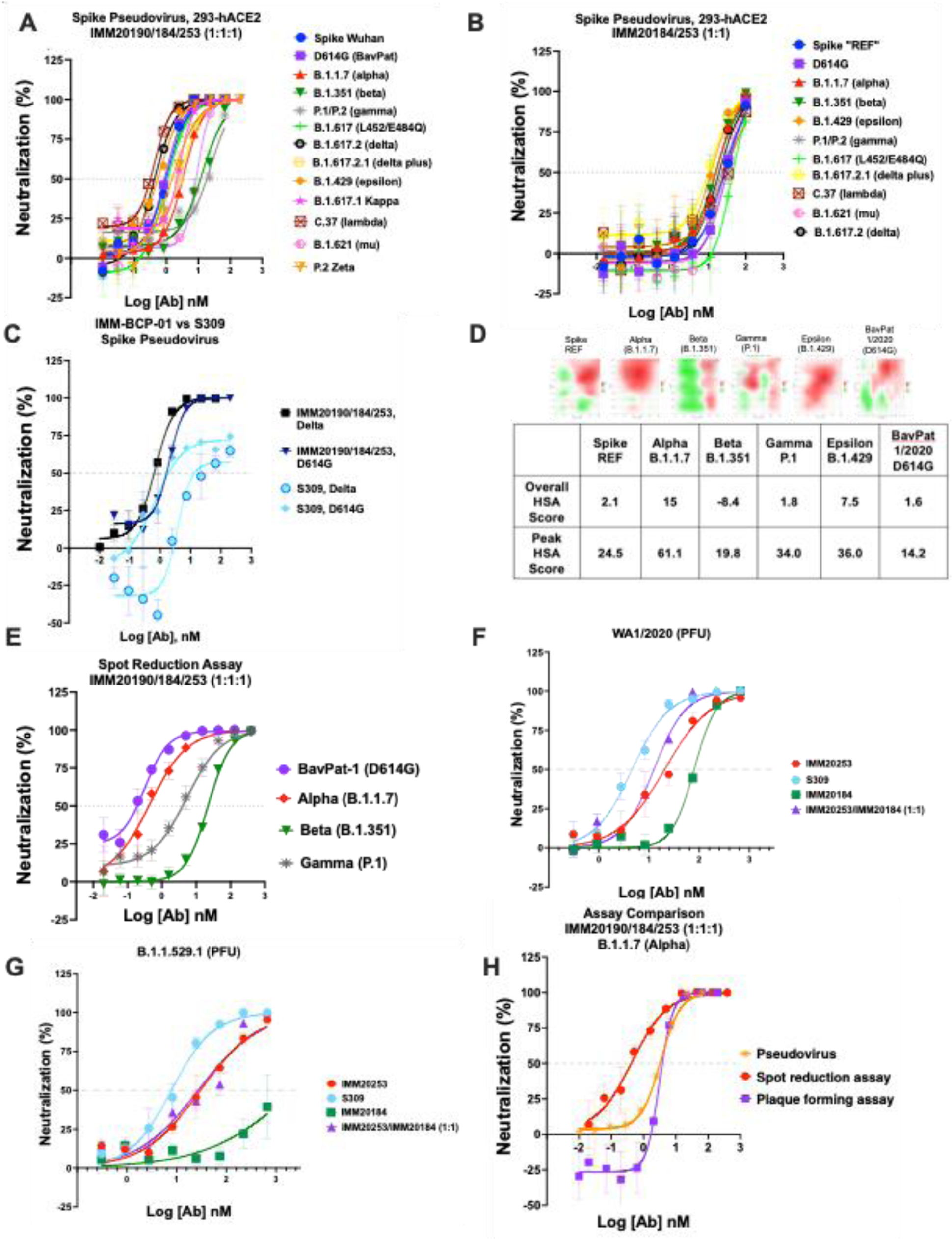
Immunome antibody combination has a synergistic effect against clinically relevant SARS-CoV-2 variants. (A) IMM20190/184/253 and (B) IMM20185/253 combination neutralizes Spike pseudoviruses that correspond to CDC VBMs. (C) Neutralization of D614G and Delta Spike pseudoviruses by three antibody cocktail IMM20190/184/253 and S309 antibodies. Shown data are representative experiments of two independent repeats. (F) Synergy scores of the 3-Ab cocktail IMM20190/184/253 against 5 pseudoviruses and one BavPat1/2020 (D614G) live virus isolate calculated with SynergyFinder 2.0 online tool. HSA, the <u>h</u>ighest <u>s</u>ingle <u>a</u>gent model score, calculates the excess over the maximum single antibody response. HSA score below -10 indicates competition; between -10 to 10 shows additive effect; and above 10 demonstrates synergy among tested agents. (E) Neutralization of VOI/VOC isolates of SARS-CoV-2 by IMM20190/184/253 cocktail as measured by ViroSpot assay. Antibodies are mixed at equimolar ratio. (F) Plaque reduction assay of REF (WA1/2020) and (E) Omicron (BA.1) virus isolates in the presence of IMM20184, IMM20253, IMM20184/253 combination and S309 antibody. (H) Neutralization of Alpha (B.1.1.7) variant by IMM20190/184/253 cocktail measured with three different methods, including a pseudovirus neutralization, intact virus spot reduction (ViroSpot) and intact virus plaque formation assays.

**Figure 4.**
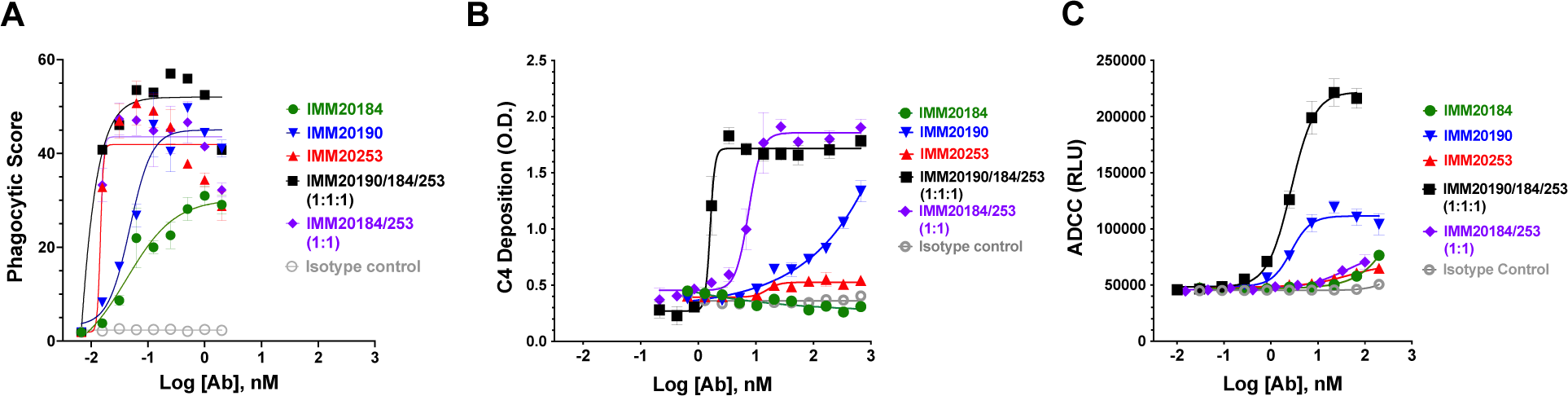
Three antibody cocktail activates potent effector function responses *in vitro.* (A) Opsonization with single IMM antibodies and two antibody (IMM20184/20253) and three-antibody (IMM190/184/253) combinations induce phagocytosis of Trimer-coated beads. (B) Deposition of classical complement component C4 on IMM antibodies bound to Trimer-coated surface.(C) Activation of antibody-mediated cellular cytotoxicity (ADCC) by IMM antibodies and two antibody (IMM20184/20253) and three-antibody (IMM190/184/253) combinations bound to S-expressing cells. Denoted points are Mean; error bars are SEM.

**Figure 5.**
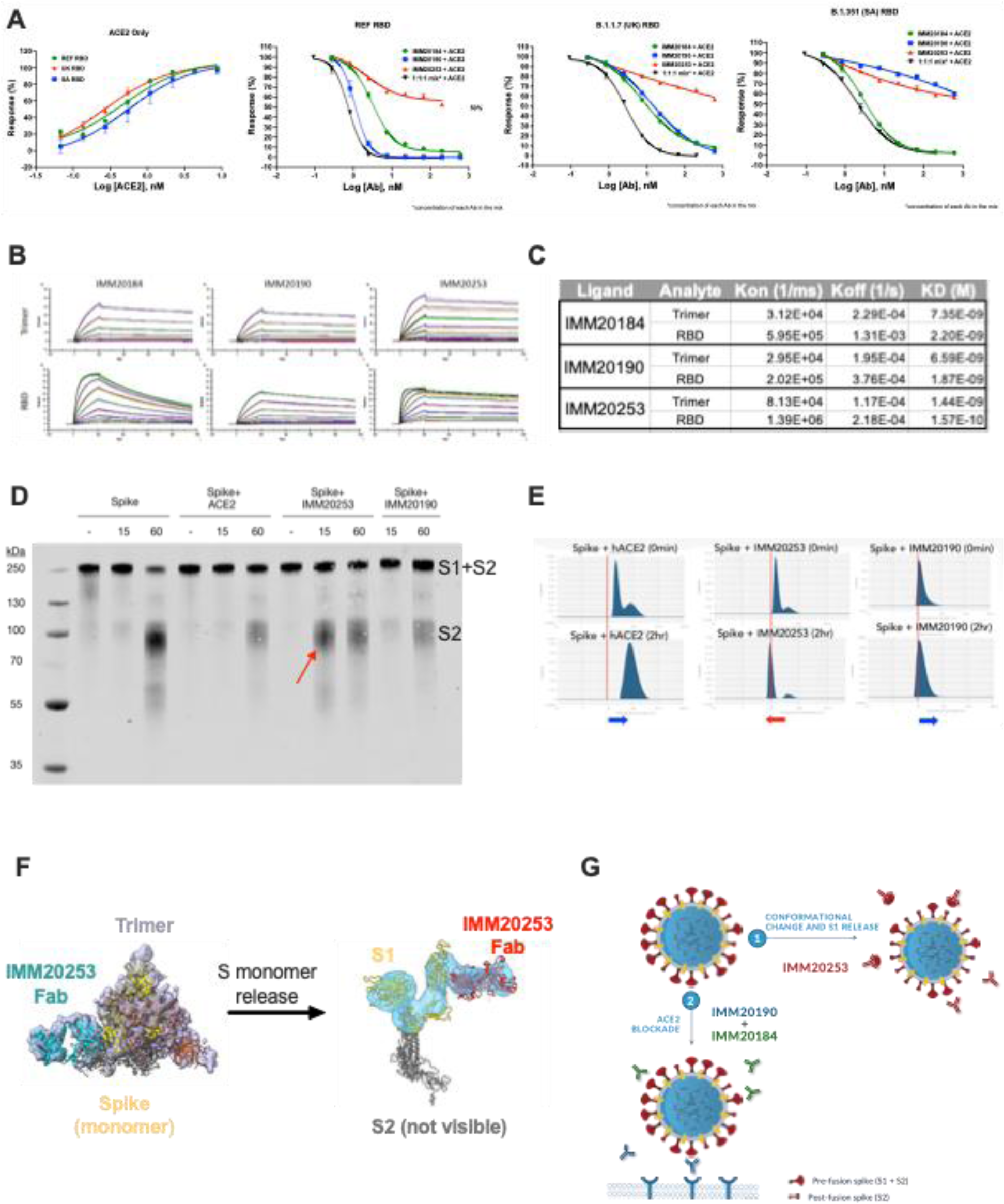
IMM20253 antibody inhibits virus in non-ACE2 dependent manner and facilitates the release of S1 protein. (A) 3-Ab cocktail inhibits RBD binding to its cellular receptor ACE2. ELISA-based receptor competition assay. (B). Antibody binding kinetics of IMM20184, IMM20190 and IMM20253 antibodies to soluble RBD and Trimer (REF variant) measured using Surface Plasmon Resonance (SPR). (C) KD, Kon and Koff values of IMM20184, IMM20190 and IMM20253 antibodies measured using a multi-cycle kinetics protocol assuming 1:1 interaction model. (D). Western blot analysis of Trimer digested with protease K after 0, 15 and 60 min in the presence of either human ACE2, IMM20253 or IMM20190. Anti-S2 staining reveals S monomer (S1+S2) and S2 protein. (E) Dynamic light scattering (DLS) analysis of Trimer complex with either ACE2, IMM20253 or IMM20190 immediately or after 2 hours incubation. (F) IMM20253 Fab binding to Trimer triggers complex disruption and release of S monomers. (G) Schematic of mechanism of action of IMM20184/190/253 or IMM-BCP-01 cocktail.

Structural analysis of an S protein timer (Trimer) complexed with bound Fabs of IMM20184, IMM20190 or IMM20253 (**Fig 1A**) identified binding patterns of IMM-BCP-01 antibodies. A final 3D reconstruction of cryo-electron microscopy (cryo-EM) micrographs of IMM20184 Fabs bound to Trimer revealed a 3:1 (Fab:Trimer) complex at 7 Å resolution with a decreased density in the Trimer core indicating disruption of the Trimer into S protein monomers along with a large reorganization of the RBD domains (**Fig 1A** **and Supp.** **Fig 1A, B**). Cryo-EM micrographs of IMM20190 Fab complexed with Trimer revealed a 3:1 (Fab:Trimer) complex at 6 Å (**Fig 1A, Supp** **Fig 1A,B**). While the variable regions of IMM20190 Fabs were clearly resolved, the constant regions were scattered, suggesting a dynamic binding nature of this antibody. Finally, the cryo-EM analysis of IMM20253 Fab-Trimer complex was repeated twice with 3:1 and 6:1 molar ratios (Fab:Trimer), with the same unexpected conclusion. The samples were not aggregated, observed with good contrast, and clearly converged into two structural families (**Fig 1A** **and Supp.** **Figure 2A, B**). The first family consisted of 1 Fab:1 Trimer complex that had one S monomer partially unfolded (revealed by a lower density). The second family included smaller complexes that converged into a 3D structure of IMM20253 Fab bound to S1 (**Fig 1A**, *side view*, and **Supp.** **Fig 1A**). The S2 portion of the spike monomer was not visible in the density maps, suggesting it moves freely in the complex relative to the S1 domain.

The Trimer reorganization induced by both IMM20184 and IMM20253 Fabs prompted us to determine their epitopes at higher resolution. Cryo-EM structures of RBD with simultaneously bound to Fabs of both IMM20184 and IMM20253 were resolved to ∼3.9 Å (**Fig 1B, Supp.** **Fig 1D**). To achieve this resolution, ∼1.9 x 10^6^ particles were subjected to three rounds of 2D classification analysis, ∼6 x 10^5^ particles were selected for ab initio reconstruction and ∼1.7 x 10^5^ particles were used for the final 3D reconstruction at a nominal resolution of 3.87 Å (**Fig 1B** and **Supp.** **Fig 1D**). The complex structure demonstrated that both IMM20184 and IMM20253 Fabs simultaneously bound to RBD protein. Consistent with **Fig 1A**, the epitope of IMM20253 was located on the outer surface of the RBD, whereas the epitope of IMM20184 faced inward and sideways, potentially enablling avid binding of IMM20184. Of note, binning of IMM20184/190/253 antibodies using bio-layer interferometry (BLI) confirmed the cryo-EM data and showed the antibodies do not compete for binding of S (**Supp.** **Fig 1H**).

The structural data was further confirmed through use of an alanine-scanning shotgun mutagenesis approach (*20*) (**Fig 1C**). In brief, we used a validated library of RBD (Wuhan) proteins expressed on the surface of HEK-293T cells, each containing one amino acid mutation (*20*). Consistent with cryo-EM data (**Fig 1A,B** **and Supp.** **Fig 1**), mutagenesis identified unique, non-overlapping epitopes for the three antibodies (**Fig 1C**). IMM20184 bound to a highly conserved region in the core RBD (**Fig 1**). The binding site lays in close proximity to the previously reported epitopes of CR3022 and COVA1-16 antibodies that bind to a cryptic epitope on RBD (*21, 22*). The IMM20184 epitope includes residues N370, F374, K378, and SP383-384 that are completely conserved among all current and previous SARS-CoV-2 VOC (**Fig 1E**), including Omicron and Delta variants.

IMM20190 bound to an epitope that included the receptor-binding ridge and an area adjacent to the receptor-binding loop. The epitope mapping analysis identified 10 residues in the RBD that interacted with IMM20190. Of these, two residues, K417 and N501 are mutated, either singly (K417 in Alpha/B.1.1.7) or doubly (K417/N501 in Beta, Gamma, or Omicron) in prior and present VOC (**Fig 1E**). The Delta variant and WA1/2020 reference strain are conserved at all 10 interaction residues. The broad epitope may explain the resistance of IMM20190 to the majority of single- and double-point mutations within the RBD region (**Supp. Table 1**) and the flexible nature of Fab binding observed by Cryo-EM (**Fig 1A**).

**Table 1.**
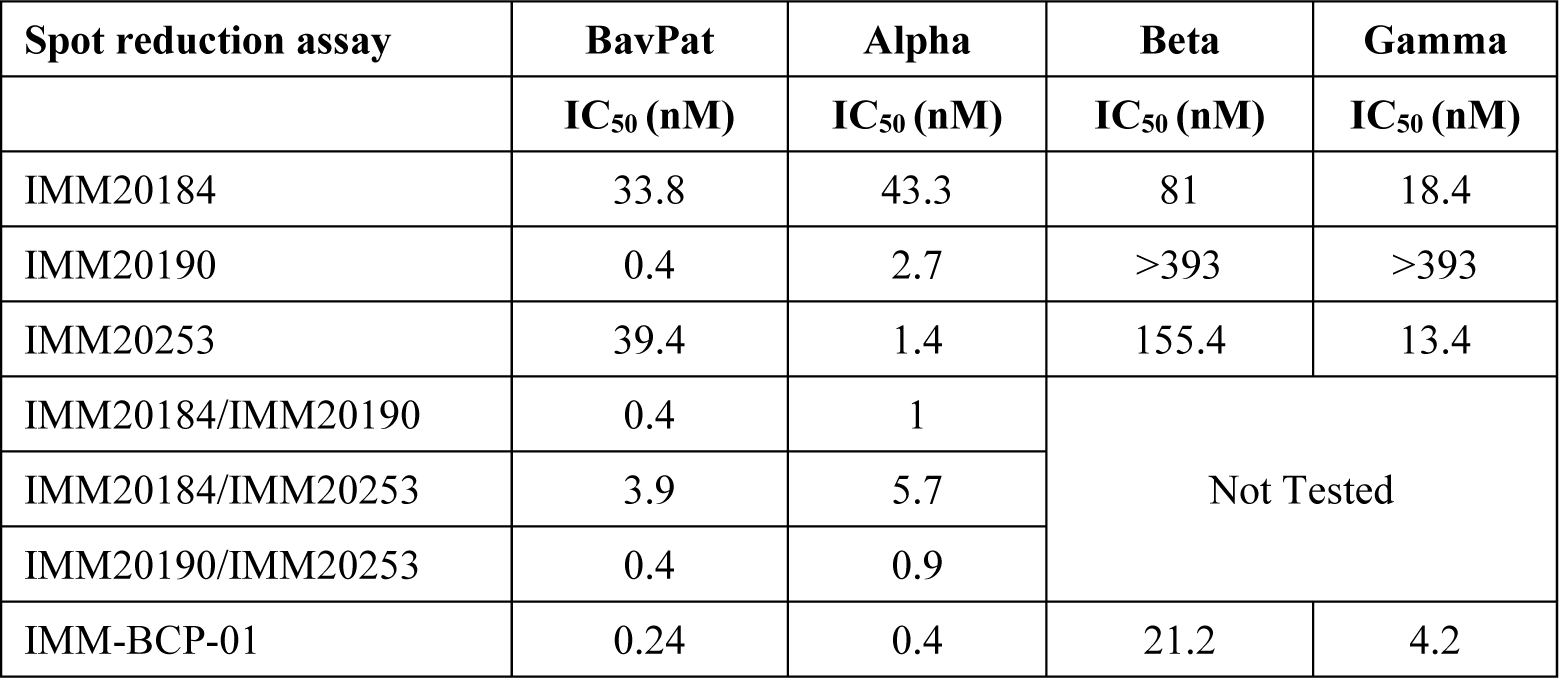
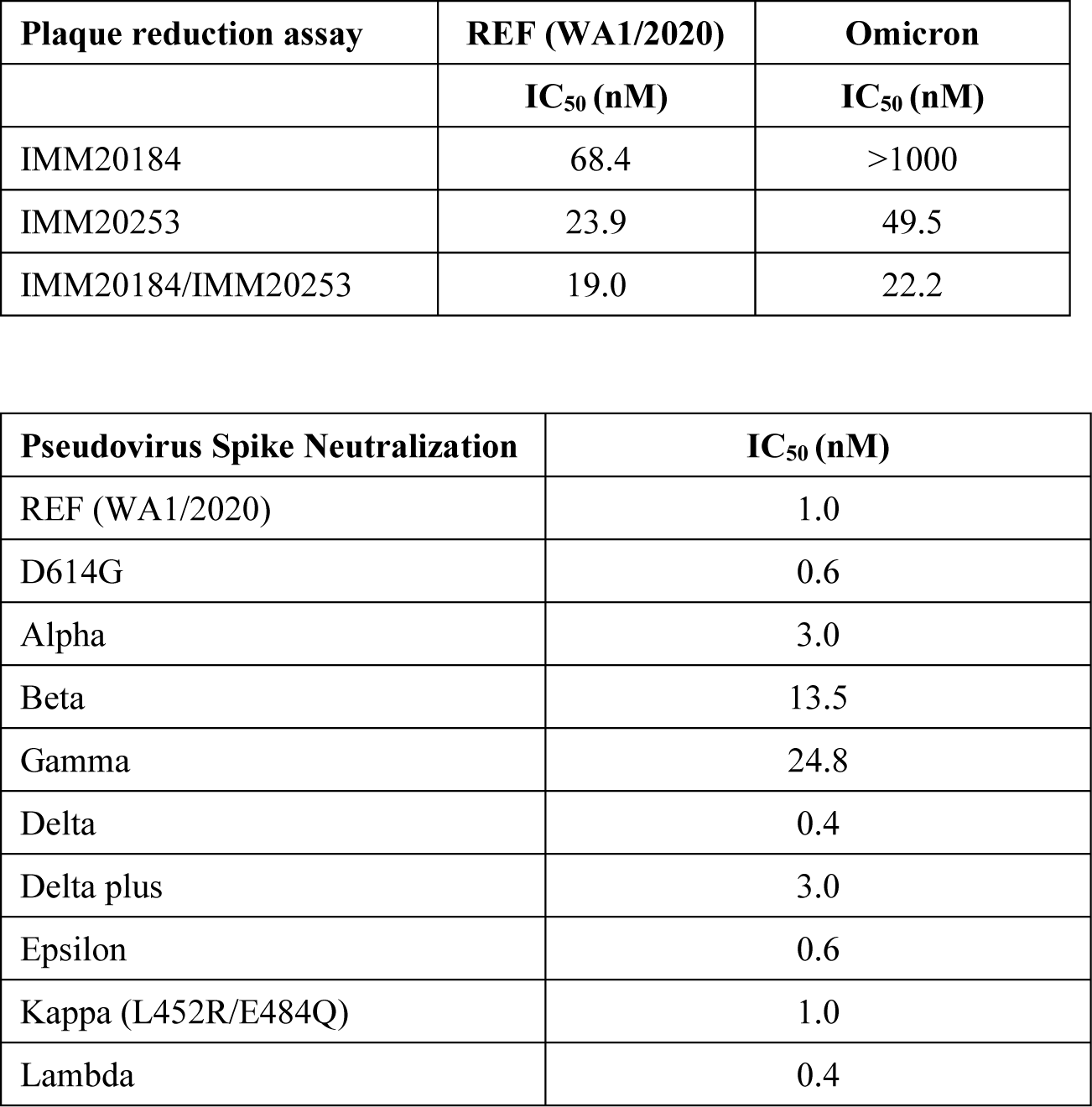
IC50 Values (nM) for IMM20184/190/253 antibodies and its combinations against SARS-CoV-2 variants. Shown values are Means generated using three different methods, including a live (authentic) virus spot reduction assay, a live (authentic) virus plaque reduction assay and a pseudovirus reporter neutralization assay.

| Spot reduction assay | BavPat | Alpha | Beta | Gamma |
| --- | --- | --- | --- | --- |
|  | IC<sub>50</sub> (nM) | IC<sub>50</sub> (nM) | IC<sub>50</sub> (nM) | IC<sub>50</sub> (nM) |
| IMM20184 | 33.8 | 43.3 | 81 | 18.4 |
| IMM20190 | 0.4 | 2.7 | >393 | >393 |
| IMM20253 | 39.4 | 1.4 | 155.4 | 13.4 |
| IMM20184/IMM20190 | 0.4 | 1 | Not Tested |  |
| IMM20184/IMM20253 | 3.9 | 5.7 |  |  |
| IMM20190/IMM20253 | 0.4 | 0.9 |  |  |
| IMM-BCP-01 | 0.24 | 0.4 | 21.2 | 4.2 |

| **Plaque reduction assay** | **REF (WA1/2020)** | **Omicron** |
| --- | --- | --- |
|  | **IC<sub>50</sub> (nM)** | **IC<sub>50</sub> (nM)** |
| IMM20184 | 68.4 | >1000 |
| IMM20253 | 23.9 | 49.5 |
| IMM20184/IMM20253 | 19.0 | 22.2 |

| **Pseudovirus Spike Neutralization** | **IC<sub>50</sub> (nM)** |
| --- | --- |
| REF (WA1/2020) | 1.0 |
| D614G | 0.6 |
| Alpha | 3.0 |
| Beta | 13.5 |
| Gamma | 24.8 |
| Delta | 0.4 |
| Delta plus | 3.0 |
| Epsilon | 0.6 |
| Kappa (L452R/E484Q) | 1.0 |
| Lambda | 0.4 |

Alanine scanning mutagenesis identified only two critical residues for IMM20253 binding to RBD, K356 and R466, located on the outer surface of the RBD. This complements the cryo-EM data (**Fig 1A, B**). K356 resides within the surface area buried by the VL, and R466 resides within the surface area buried by the VH of IMM20253. R466 residue is conserved in all sarbecoviruses or lineage b betacoronaviruses, whereas K356 is conserved in most ((**Fig 1E**) and (*21, 23, 24*)). In summary, IMM20253 binds to a highly conserved epitope on the outer surface of RBD, does not compete with IMM20184 and IMM20190 mAbs for S binding and induces dissociation of Trimer complex into monomers.

### IMM-BCP-01 cocktail efficiently suppresses the severity of the disease in an *in vivo* model of SARS-CoV-2 infection

We tested the efficacy of different combinations and doses of these three antibodies in Syrian Golden hamsters inoculated with SARS-CoV-2 (WA_CDC-WA1/2020) (**Fig 2A**). When administered to animals 6 hours after viral challenge (treatment paradigm), we observed that IMM20190 or 2-Ab combinations of IMM20184/IMM20190 or IMM20190/IMM20253 led to robust viral clearance (**Fig 2B**). However, the greatest clearance was observed with the 3-Ab cocktail. Five of the six animals in this cohort showed an approximately 2.5-log10 reduction of viral titer in the lung on day 4 post viral challenge. In a follow-up study, the 3-Ab cocktail decreased the viral titer in the lungs of animals inoculated with a high-titer (3.3 x 10^5^ TCID50) of WA_CDC-WA1/2020 (REF variant) by over 100-fold. This efficacy was observed when the antibodies were administered at either 1:1:1 (p=0.0077) or 1:0.5:0.5 (IMM20190:IMM20184:IMM20253; p=0.0143) molar ratios (**Fig 2C**). However, when subjected to an F-test, the variability in clearance level in 1:0.5:0.5 group was higher (P < 0.0001) than when animals were treated with an equimolar ratio cocktail (**Fig 2C**). These studies were performed using a viral inoculum that was approximately 10-fold higher than what is typically used to evaluate efficacy of antibody therapies (*23, 25*). When repeated at a lower inoculating dose (3.3 x 10^4^ TCID_50_ per animal) (**Fig 2D**), treatment of hamsters with IMM-BCP-01 (0.1 mg each, 0.3 mg total antibody) resulted in a significant (p<0.0080) ∼3.5 log10 decrease in viral titer relative to vehicle-treated controls. Taken together, these data support the IMM-BCP-01 cocktail as comprising all three antibodies at 1:1:1 ratios to obtain the most consistent level of viral clearance.

### IMM20184, IMM20190 and IMM20253 antibody combinations demonstrates a dose-dependent inhibition of virus load in lungs of hamsters infected with WA1/2020, Alpha, Beta and Omicron variants of SARS-CoV-2

IMM-BCP-01 cocktail was designed to recognize and inhibit variants that have and could emerge. Consistent with that goal, IMM-BCP-01 exhibited a dose-dependent inhibition of all viral variants tested *in vivo*, including the reference (WA1/2020) variant, Alpha, Beta and Omicron isolates (**Fig 2E, F**). The 3-Ab cocktail suppressed viral infection in the lungs of hamsters pre-treated with doses as low as 0.1 mg of each antibody (0.3 mg total dose) 24 hours prior to virus challenge. Higher doses of IMM-BCP-01 lowered viral loads in hamsters to a greater degree. A 10,000-fold reduction in viral load in lungs of animals inoculated with WA1/2020, and a 1,000-fold reduction in animals inoculated with Alpha and Beta isolates were achieved with doses of 0.3 and 0.5 mg each (0.9 mg and 1.5 mg total for a IMM20190/184/253 cocktail). Animals infected with Omicron isolate developed a lower viral lung load (∼4.4*10E4 PFU/g) comparing to other isolates, that was dose-dependently reduced by a standalone IMM20253 antibody. 2-Ab combination IMM20153/IMM20184 (0.5 mg ea or 1 mg total dose) further decreased viral load in lungs to levels comparable to the lower limit of detection (LOD) for the study. Thus, IMM20184, IMM20190 and IMM20253 antibody combinations potently suppresses infection of multiple SARS-CoV-2 variants *in vivo* in a dose-dependent manner.

### IMM-BCP-01 cocktail exposure and pharmacokinetics

When administered via i.p. injection, the IMM-BCP-01 cocktail generally followed first-order absorption and elimination process with a half-life of approximately 100 hours in hamsters (**Supplemental Fig 2A, B**). Unexpectedly, we observed that some animals treated with IMM-BCP-01 had lung titers equivalent to non-treated controls. To better understand that lack of effect, terminal bleeds were assessed for levels of human IgG in the plasma. Those studies demonstrated that variability in viral clearance correlated directly with systemic distribution of IMM-BCP-01 (**Supplemental Fig 2C**). Animals that exhibited lower viral lung titers were associated with terminal plasma levels of IMM-BCP-01 greater than 3-5 µg/mL. In contrast, IMM-BCP-01 was not observed at appreciable levels in the blood of animals that failed to clear virus from the lungs, that rather reflects the difficulties with antibody injection to these animals. Effective levels of IgG in the blood were achieved with dose levels as low as 0.1 mg each (0.3 mg total dose) in both the prophylactic and treatment settings when the drug was absorbed and systemic exposure was achieved (**Suppl. Fig 2C**).

### IMM-BCP-01 has a combinatorial neutralizing effect against current and prior VOCs of SARS-CoV-2 virus

IMM-BCP-01 cocktail was evaluated in three live (authentic) virus neutralization assays and one reporter pseudovirus assay (**Fig 3**) using an array of viral variants. The three independent ive virus neutralization assays provided comparable data, which agreed with pseudovirus neutralization tests (**Table 1**). The antibody cocktail neutralized all tested VBM and VOC (**Fig 3**). The IMM-BCP-01 cocktail (IMM20184/190/253), as well as IMM20184/20253 combination, completely neutralized all pseudovirus variants tested (**Fig 3A,B**). Overall, IMM-BCP-01 potently neutralized the spectrum of variants tested, with all IC50 values being within 2-log of the reference pseudovirus encoding a WA1/2020 S protein. The 3-Ab cocktail had a modest, but reproducible, *increase* in potency against Delta, Lambda (C. 37), and Epsilon (B.1.429) pseudoviruses, which could be explained by a higher susceptibility of Trimers from these variants to structural rearrangements. In context of current landscape of antibody therapeutics for COVID-19, IMM-BCP-01 outperformed S309 against Delta and a WA1/2020 D614G pseudovirus (**Fig 3C**). S309 is the parental clone of VIR-7831, which obtained an EUA and retains activity against some Omicron variants (*25*).

To better understand the 3-Ab cocktail, we performed a series of experiments focusing on the combinatorial contributions of the component antibodies to the overall neutralizing activity. We tested 2-Ab mixtures of IMM20190 (1x concentration) with either IMM20184 (1x) or IMM20253 (1x), and a 3-Ab combination of IMM20190 (1x) with IMM20184/IMM202053 (0.5x each) in pseudovirus neutralization assays (**Supp.** **Fig 3**). Double and triple antibody combinations dose-dependently neutralized pseudovirus variants corresponding to three VOC (**Supp.** **Fig 3A, B)**. We calculated each antibody contribution to the observed neutralizing effect using SynergyFinder 2.0 (*26*)A score below -10 suggests an antagonistic (competitive) effect; a score between -10 and 10 reflects an additive effect; and a score above 10 suggests a synergistic effect of the combined treatment. We detected a concentration-dependent synergistic potential of combinations (**Supp.** **Fig 3C**). In variants that IMM20190 potently neutralized, such as WA1/2020 and Epsilon, antibody combinations are mainly additive, as IMM20190 neutralization was sufficient and did not require the two other antibodies. In variants where the potency of IMM20190 was reduced, such as Beta and Gamma, combinations were also additive. In addition, IMM20184 and IMM20253 antibodies as a double combination had an additive neutralizing effect against these variants (**Supp.** **Fig 3D**). We observed the highest synergistic potential of the 3-Ab combination IMM20190/184/253 for the Alpha variant, where each of the single antibodies neutralized with comparable IC50’s (Fig 2D). In this variant background, the triple antibody combination outperformed all three of the individual antibodies. Thus, the 3-Ab cocktail neutralized all tested variants and was associated with additive or synergistic effects depending on the strain. Combined with the observed antibody pharmacokinetics data (**Supp.** **Fig 2A**), these data suggest that administration of the 3-Ab cocktail (0.5 mg each) reaches serum concentrations in vast excess of the IC50 neutralization concentrations observed for all SARS-CoV-2 variants tested, including the Alpha, Beta, Gamma, and Delta (**Supp.** **Fig 2, and Fig 2, 3**).

When extended to intact virus isolates, we observed equivalent, or better potency of the IMM-BCP-01 cocktail against WA1/2020, BavPat (D614G), Alpha, Beta, and Gamma variants measured in focus (**Fig 3E**) or plaque (**Fig 3F, G**) reduction assays as compared to the corresponding pseudovirus neutralization assay (**Table 1**).

IMM-BCP-01 was evaluated for activity against two different live virus isolates of the Omicron BA.1 variant, as well as an Omicron variant, BA.1.1, harboring an additional R346K mutation (**Fig 3F, G; Supp.** **Fig 4**). Consistently with the data observed *in vivo* (**Fig. 2H**), a standalone IMM20253 antibody neutralized Omicron (BA.1) authentic virus in plaque reduction assay (Fig 3H) and BA.1 and BA.1.1 in focus reduction neutralization assays (**Supp.** **Fig 4**). Although no mutations present in the Omicron isolates mapped to critical binding residues for IMM20184 (**Fig 1C**), the antibody lost neutralization potency. A partial loss of IMM20184 activity was observed in plaque reduction assays with BA.1, but complete loss of neutralizing activity was observed against BA.1.1 in the context of the FRNT assay (**Fig 3G, Supp. Fig 4**). The IMM20184/253 combination showed an additive effect, compared to the IMM20253 antibody alone, in the plaque assay (**Table 1**) and (Fig. 3G), that is in agreement with the result observed using Omicron isolate *in vivo* (**Fig. 2H**).

Finally, we observed a higher *in vivo* potency of IMM-BCP-01 cocktail, compared to its activity by virus neutralization assays *in vitro*. We detected a 100-fold increase in EC50 of IMM-BCP-01 cocktail against Beta variant in both pseudovirus (**Fig 3A**) and authentic virus (**Fig 3E**) assays *in vitro* that only resulted in a minor dose increase (from 0.3 mg to 0.5 mg per antibody) *in vivo* (**Fig 2F, G and 3A**), outlining the importance of *in vivo* studies for anti-SARS-COV-2 antibodies. We performed *in vitro* neutralization assays at multiple facilities, including academic and industry laboratories, and observed a ∼10-fold difference in EC50s values for same virus variants (Alpha variant as an example, **Fig 3H**).

### IMM20190/184/253 antibody cocktail activates potent effector responses *in vitro*

A growing body of evidence suggests that intact effector functions are required for optimal viral clearance in animal models of COVID-19 (*27–29*). The antibodies comprising IMM-BCP-01 retain intact IgG1 Fc domains and bind to the RBD in a non-competitive manner (**Fig 1, Supp.** **Fig 1H**). We hypothesized that IMM2019/184/253 might generate an oligoclonal response to S protein that activates Fc-mediated effector functions including antibody-dependent cellular cytotoxicity (ADCC), antibody-dependent cellular phagocytosis (ADCP), and classical complement pathway (CP) (Fig 4). To test this hypothesis, we first measured antibody-induced phagocytosis of Trimer-coated beads using a published method (*30*). All three human antibodies induced phagocytosis of Trimer-coated beads in a dose-dependent manner relative to an IgG1 isotype control (**Fig 4A**). Even a low (∼15 pM) concentration of IMM20253 antibody potently induced phagocytosis. The 3-Ab cocktail (IMM20190/184/253) demonstrated a higher phagocytic score than a 2-Ab cocktail (IMM20184/253) or each individual antibody (**Fig 3A**). We did not observe phagocytosis of Trimer-coated beads in the presence of an IgG1 isotype control antibody. Next, we evaluated activation of the classical CP by IMM20190/184/253 cocktail (Figure 3B). In brief, we adapted a CP activation assay (*31*) and measured deposition of the complement component C4 from serum of normal human donors on anti-S antibodies bound to Trimer-coated surface. IMM20190 and IMM20253 antibodies bound to Trimer promoted detectable levels of C4 deposition. While IMM20184 binding to Trimer alone did not activate CP in this assay, the 2-Ab cocktail IMM20184/253 induced C4 deposition on antibody-Trimer complexes (**Fig 3B**). The three antibody cocktail IMM20190/184/253 induced the most robust activation of C4 deposition. Since all tested antibodies had the same intact heavy chain IgG1 Fc region, we hypothesized that C4 deposition on Trimer-Ab immune complex might depend on the Fc epitope conformation and accessibility as previously demonstrated for other antibodies (*32, 33*). Cryo-EM studies (**Fig 1, Supp.** **Fig 1**) indeed demonstrated that IMM-BCP-01 cocktail attacks Trimer from different directions and may indeed create an array of Fc regions that facilitates binding of C1q. Finally, ADCC assay revealed a similar synergy among IMM20184/190/253 antibodies (Fig 3C). While each antibody induced a mild (IMM20184, IMM20253) to moderate (IMM20190) activation of ADCC, the three antibody cocktail IMM20190/184/253 induced the greatest response (**Fig 3C**).

### IMM20184, IMM20190, but not IMM20253, block S interactions with ACE2

The location of IMM20184 and IMM20190 epitopes suggests that these two antibodies likely block ACE2 binding. To test this hypothesis, we performed a biochemical ELISA-based receptor inhibition assay. The affinity of a soluble ACE2 protein to Wuhan-1, Alpha, and Beta variant RBDs coated to an ELISA plate was evaluated in the presence of various concentrations of antibodies of interest. Consistent with the data from the homogeneous time resolved fluorescence (hTRF) assay (**Supp. Table 1**), IMM20184 potently inhibited ACE2 binding to all three RBD variants (**Fig 5A**). IMM20190 blocked ACE2 binding to Wuhan-1 and Alpha variant RBD proteins, and partially decreased ACE2 binding to the Beta variant RBD. In contrast, IMM20253 did not fully block ACE2 interactions with any of the three RBD variants tested (**Fig 5A**). A partial inhibition of ACE2 binding by IMM20253 (up to 40%, depending on the concentration) was detected for all three S variants tested. These data are consistent with the location of the IMM20253 epitope relative to the ACE2 binding site and suggest its neutralization occurs through a distinct mechanism of action (see below). Finally, an equimolar mixture of IMM20190, IMM20185, and IMM20253 antibodies disrupted ACE2 binding to all three tested RBD variants. The inhibitory effect of a 3-Ab cocktail was more pronounced than the effect of each individual antibody. Of note, we detected a minor ACE2 binding preference to Alpha RBD than to Beta RBD variant (**Fig 6A**, *left panel*).

### IMM20190, IMM20184 and IMM20253 bind to soluble RBD and S1 proteins recapitulating different SARS-CoV-2 variants

Steady-state binding of the three antibodies across a range of variants was characterized via hTRF (**Supp. Table 1 and Supp. Fig 6**). Each of the three antibodies was tested for binding against isolated RBD or S1 protein encompassing over 20 different single and multiple mutations that correspond to naturally occurring and predicted escape mutations. IMM20184 and IMM20253 retained picomolar EC50 binding to most of single and multiple point mutations tested, including those present in VBM and VOC. Furthermore, IMM20253 exhibited some binding to SARS-CoV-1 Spike. In contrast, IMM20190 binding was reduced, to varying degrees, by K417N and the series of RBD-localized mutations associated with K417N/E484K/N501Y or K417T/E484K/N501Y variants. Binding of IMM20190 appeared to be partially restored for two different S variants containing D614G even in the presence of K417N (**Supp. Table 1**).

### Analysis of binding kinetics of IMM20184, IMM20190, and IMM20253

Antibody binding kinetics were measured using a multi-cycle kinetics protocol on a Biacore T200 surface plasmon resonance instrument. All three antibodies bound with high affinity to both RBD and Trimer. (**Fig 5B, C**). All three antibodies bound with rapid on-rates (ka > 2.95+E4 1/Ms) to Trimer, but exhibited even faster on-rates (ka) to RBD. This effect was least pronounced for IMM20190, suggesting that its epitope is the most accessible in the intact Trimer structure. IMM20184 and IMM20253 bound to the RBD 17- and 19-fold faster than to the Trimer, respectively. The dissociation of IMM20184 from Trimer (k ^Trimer^ = 2.3E-04 1/s) was 6-fold slower than from RBD (k ^RBD^ =1.3E-03 1/s). This difference in dissociation suggests the antibody/Trimer complex is stabilized through an avid binding mechanism between the antibody and two subunits of the Trimer, consistent with the epitope mapping data (**Fig 1**). IMM20253, despite exhibiting the greatest difference (19-fold) in on-rates between the RBD and Trimer, has the fastest on-rate for the Trimer of all three antibodies. This result suggests the epitope is readily accessible in the context of the Trimer structure.

### IMM20253 binding releases Spike monomers and facilitates protease cleavage

IMM20253 disruption of Trimer into Spike monomers, detected by cryo-EM (**Fig 1**), suggested that the antibody might facilitate cleavage of S into S1 and S2. We used a previously published method to evaluate Trimer sensitivity to protease cleavage in the presence of an antibody (*34*). Briefly, Trimer protein was mixed with protease K in the presence of either (1) buffer only, (2) human recombinant ACE2 protein, (3) IMM20253 or (4) IMM20190, an anti-S antibody that recognizes ACE2-binding region, and incubated for 0, 15 and 60 minutes. Protease readily cleaved S incubated with buffer after 60 min. ACE2 or IMM20190 appeared to partially decrease protease cleavage at 60 min, perhaps due to steric hindrance. In contrast, IMM20253 induced S cleavage after 15 min (**Fig 5D**). In a complementary experiment, we evaluated S samples preincubated with ACE2, IMM20253 or IMM202190 in the absence of protease. Samples were analyzed using a standard “premix” protocol by dynamic light scattering (DLS). Incubation of S with ACE2 or IMM20190 led to the generation of complexes and increased the size of particles, measured as an increase in hydrodynamic diameter after 2 hours of incubation (Figure 5E). Consistent with cryo-EM and protease cleavage results, incubation of S with IMM20253 decreased the hydrodynamic diameter of the resulting complexes, consistent with complex disruption. These data support a unique mechanism of action for IMM20253 whereby binding disrupts the Trimeric architecture of the S complex and facilitates cleavage into S1 and S2 in the presence of proteases.

## DISCUSSION

We have described three unique antibodies with different mechanisms of action that bind to non-competing epitopes on S protein, trigger Trimer reorganization into one resembling a post-fusion confirmation, and induce a potent antiviral response *in vitro* and *in vivo*. Further, we revealed a unique mechanism of action of IMM20253; this antibody binds to a conserved epitope on Trimer Spike and induces complex dissociation into monomers and cleavage into S1 and S2 subunits. When combined with IMM20190 and IMM20184, the 3-Ab cocktail consistently showed a robust antiviral potency *in vivo* and *in vitro*, neutralized all VOC and VBM tested and induced a potent multicomponent Fc effector response.

We previously reported the generation of several hundred of anti-SARS-CoV-2 antibodies with diverse antiviral properties (*19*). Subsequent functional analysis guided us to the selection of three unique, non-competitive anti-S antibodies with potent yet complementary antiviral effects that bound to three spatially distinct surfaces of RBD region of Spike protein. While each antibody was capable of neutralizing many viral variants, the antibody cocktail robustly neutralized all variants tested with additive or synergistic effect.

Each of the three antibodies comprising IMM-BCP-01 appear to elicit viral neutralization through different mechanisms. IMM20190 and IMM20184 antibodies compete with a cellular receptor hACE2 for binding to Spike protein. The IMM20190 epitope, identified by Cryo-EM and confirmed using mutagenesis, extends to two surfaces, including the receptor binding ridge of RBD and the region around the receptor-binding loop (*35*). Considering the breadth of this epitope, IMM20190 was shown to be resistant to small changes in the RBD sequence. The epitope of IMM20184 antibody is located in N370 – P384 region of the RBD and consists of 5 critical residues, surrounding the RBD core (*35*), that is conserved in Omicron and all prior VOC of SARS-CoV-2 and other SARS-related coronaviruses (*21, 22*). IMM20184 antibody has a higher affinity to a soluble RBD, yet 5.7-fold slower dissociation from a soluble Trimer, indicating avid binding to Trimer. The COVA-1 antibody that bound to the area adjacent to IMM20184 epitope has been demonstrated to have strong cross-neutralizing properties due to its avid binding (*21*). The remaining antibody IMM20253 does not directly compete with hACE2, but binds to a conserved epitope (K356 and R466 residues) that is not a common target of the human immune system for generating neutralizing antibodies (*35*). IMM20253 binding leads to a dissociation of Trimer into S protein monomers, that likely facilitates cleavage into S1 and S2 in the presence of proteases. Thus, IMM20253 triggers a conformational change of S protein into its post-fusion form and prevents binding to host cells in ACE2-independent manner. IMM20253 epitope is present in all human as well as in SARS-related coronaviruses(*24*) and is, therefore, expected to be retained in emerging human SARS-related viruses. K356 has been previously shown to participate in a formation of a hydrophobic pocket in the RBD (*36*) that may explain its functional importance and evolutionary conservation. The conserved patch of amino acids around R466 has only rarely elicited an antibody response (*35*). R466 is conserved in all SARS-related pangolin and bat betacoronaviruses, making it an attractive target for the therapeutic intervention and vaccine design. There are two reported antibodies (a nanobody derived from llama and a mouse-derived antibody) that recognize larger patches of RBD and appear to have overlapping epitopes with IMM20253 (*34, 37*). However, both described antibodies bound to broader epitopes, i.e more sensitive to mutational drift; both were generated via animal immunizations and need to be tested for off-target binding to human tissues; and either need to be humanized (mouse-derived) or re-designed (llama-derived) prior to consideration as therapeutics. These two studies, however, further confirm the importance of the IMM20253 epitope.

Importantly, the range of neutralization potency exhibited by IMM-BCP-01, across the breadth of pseudovirus and live virus tested, translated into *in vivo* efficacy in animal models. This was illustrated most notably in the setting of Beta and Omicron variants. Despite showing a variable level of *in vitro* neutralization potency (4.9 – 24 nM) against the Beta isolate depending on the assay used to measure activity, IMM-BCP-01 exhibited robust *in vivo* efficacy at doses consistent with those currently being used in the clinic for other SARS-CoV-2 antibodies. While IMM-BCP-01 appears to neutralize virus comparably to S309, the *in vivo* potency of IMM-BCP-01 exceeds that of VIR-7831 (*18*). Consistent with the results obtained for Beta variant, a modest neutralization of Omicron variant by IMM20253 alone (49.5 nM) or by IMM20184/20253 combination (22 nM) *in vitro* translated into a striking potency of IMM20184/IMM20253 combination against Omicron variant *in vivo*, decreasing the virus lung load in Omicron infected hamsters ∼100 fold to the level comparable to lower limit of detection for the method. We hypothesize that neutralization potency alone does not account for the overall potency *in vivo* as compared to what was observed for in the published literature. Published data for REGN10933 and REGN10987, the antibodies comprising REGN-CoV2, suggest they were more potent in *in vitro* neutralization assays (*38*), yet did not appear to lead to higher levels of viral clearance in hamster models of COVID-19 (*39*). We demonstrate that Fc-mediated viral clearance mechanisms are also enhanced in the context of the IMM-BCP-01 cocktail as compared to any of the three individual antibodies alone. We argue that the enhanced viral clearance observed *in vivo* may be a direct result of the oligoclonal nature with which the IMM-BCP-01 antibodies bind to the RBD of S protein. The ability to neutralize via multiple mechanisms leads to synergy, in conjunction with the Fc-dependent activation of effector functions, may explain the robust potency detected in the *in vivo* experiments, and is in agreement with previous reports (*27–29*). Interestingly, no apparent clinical benefit has been derived by increasing the dose of REGN-CoV2 from 2400 mg to 8000 mg (*40*). Similarly, viral clearance of WA1/2020 elicited by VIR-7831 in hamster studies plateaued at approximately 15-fold clearance upon treatment with 5 mg/Kg of antibody; increasing to 15 mg/kg led to no better clearance (*25*). In contrast to those findings, administration of IMM-BCP-01, within the 3 – 9 mg/Kg dose range tested, yielded a dose response of ∼300-10,000-fold clearance of the WA1/2020 virus, with the 10,000-fold decrease being at the limit of detection for evaluating absolute clearance levels. Importantly, the dose response was not limited to the WA1/2020 isolate, as a similar dose-response was observed in clearance of Alpha, Beta and Omicron variants.

These three antibodies were identified using the Immunome’s Discovery Platform, based on the antibody function, target and biochemical properties. Having the structure and mechanistic data opens up the opportunity for a potential rational design of antibody combinations(*41, 42*).

In summary, we identified and characterized three potent patient-derived antibodies IMM20190, IMM20184 and IMM20253 with a unique set of antiviral properties. When combined into the IMM-BCP-01 cocktail, they synergize to neutralize multiple variants of SARS-CoV-2, potently activate Fc-mediated antiviral effector response, and demonstrate antiviral effects *in vivo*. We described IMM20253, that belongs to a unique class of human antibodies, that recognizes a rare epitope and triggers dissociation of Trimer. Based upon the data we have presented, IMM-BCP-01 is effective across the spectrum of variants known to date and should retain activity against future variants. Importantly, recent data support the idea that targeting SARS-CoV-2 with several antibodies should reduce viral escape from IMM-BCP-01 (*11*). As IMM-BCP-01 antibody cocktail may be used in both prophylaxis or therapy setting against SARS-CoV-2 variants, clinical trials are warranted.

## MATERIAL AND METHODS

### Cells

Reporter virus particles (RVP’s) were purchased from Integral Molecular, ACE2-293T cells (Integral Molecular; Cat #C-HA102) were cultured in DMEM containing 10% FBS, 10mM HEPES, and 0.5 μg/mL Puromycin. Vero E6 cells (BEI resources, NIAID, NIH: VERO C1008 (E6), African green monkey kidney, Working Cell Bank NR-596) were maintained in humidified incubators at 37°C and 5% CO2 in DMEM high glucose with GlutaMAX and sodium pyruvate (Gibco^TM^, cat #10569) and 10% certified US-origin heat-inactivated fetal bovine serum (Gibco^TM^, cat #10082). African green monkey Vero-TMPRSS2 (*43*) cells were cultured at 37°C in Dulbecco’s Modified Eagle medium (DMEM) supplemented with 10% fetal bovine serum (FBS), 10 mM HEPES pH 7.3, 1 mM sodium pyruvate, 1× non-essential amino acids, and 100 U/mL of penicillin–streptomycin with 5 μg/mL of blasticidin.

### Animal studies

All animal studies described in the manuscript were carried out under Institutional Animal Care and Use Committee (IACUC)-approved protocols at the respective institutions (BU and MRIGlobal) and where appropriate were reviewed and approved by Animal Care and Use Review Office of USAMRDC (ACURO).

### Syrian hamster model of SARS-CoV-2 infection

Syrian Golden Hamsters (Envigo) were challenged on Study Day 0 with SARS-CoV-2 via intranasal inoculation using 0.1 mL of either 1.67 × 10^5^ or 1.67 × 10^6^ TCID_50_/mL material (WA_CDC-WA1/2020) (post-exposure treatment experiment), or 1 x 10^4^PFU (WA_CDC-WA1/2020, Alpha (B.1.1.7) or Beta (B.1.351)) material (pre-exposure treatment experiment). Hamsters were either treated one day before challenge, or 6 ± 1 hour after challenge via an intraperitoneal (i.p.) injection. Animals were euthanized on Study Day 4. The lungs were harvested and homogenized for viral titer determination via TCID_50_ or plaque assay. Viral clearance levels obtained by the various treatments were compared to non-treated controls using a two-way ANOVA with Tukey’s multiple comparisons test. F-tests were performed using an online calculator (https://www.statskingdom.com/220VarF2.html).

### Pseudovirus Production and Neutralization Assay

Neutralization experiments using SARS-CoV-2 luciferase reporter virus particles (RVP’s) (Integral Molecular) were based on the manufacturer’s instructions. In brief, RVP’s were thawed for 2-3 minutes in a 37°C water bath. The recommended amount of RVP’s was added to the inner wells of a white opaque 96 well plate (Corning; Cat #3917) or 384 well plate (Greiner Bio-One; Cat #781080). Media containing the indicated amount of antibody was added to each well, resulting in a final volume of 100 μL per well (96 well plate) or 25 μL per well (384 well plate). The antibody/RVP mixture was pre-incubated for 1 hour in a 37°C incubator containing 5% CO2. ACE2-293T target cells were added to each well (2 x 10^4^ cells in 100 μL for a 96 well plate or 0.9 x 10^4^ cells for a 384 well plate) and incubated for 72 hours. Media was removed from all wells, equal volumes of PBS and Renilla-Glo Luciferase Assay Reagent (Promega; Cat #E2720) were added to each well (60 μL total for a 96 well plate or 30 μL total for a 384 well plate). After 10 minutes, luminescence was measured on the EnSpire Plate Reader (PerkinElmer). Percent neutralization was calculated with the following equation: [(RLU of Virus + cells) – (RLU of Experimental Sample)] / [(RLU of Virus + cells) – (RLU of cells only)].

### Epitope Mapping of IMM20190, IMM20184 and IMM20253 antibodies

Shotgun Mutagenesis epitope mapping services were provided by Integral Molecular (Philadelphia, PA) as described in (*44*). Briefly, a mutation library of the target protein was created by high-throughput, site-directed mutagenesis. Each residue was individually mutated to alanine, with alanine codons mutated to serine. The mutant library was arrayed in 384-well microplates and transiently transfected into HEK293T cells. Following transfection, cells were incubated with the indicated antibodies at concentrations pre-determined using an independent immunofluorescence titration curve on wild type protein. MAbs were detected using an Alexa Fluor 488-conjugated secondary antibody and mean cellular fluorescence was determined using Intellicyt iQue flow cytometry platform. Mutated residues were identified as being critical to the MAb epitope if they did not support the reactivity of the test MAb but did support the reactivity of the reference MAb. This counter-screen strategy facilitates the exclusion of mutants that are locally misfolded or that have an expression defect.

### Calculation of Synergistic Neutralization by Antibody Combinations

In the context of pseudovirus neutralization, synergy between two or three monoclonal antibodies in combination is defined as neutralization that is greater than neutralization by the most effective monoclonal antibody alone. To test whether combinations of antibodies show synergy in neutralizing SARS-CoV-2, we used an approach similar to one described previously (3). Pseudovirus neutralization experiments were set up as described above, except that multiple monoclonal antibodies were tested in combination. Briefly, for combinations of two antibodies, one test article was titrated in the background of each concentration in a serial dilution of the other test article. Single antibody titrations were included as controls. For combinations of three antibodies, one test article was titrated in the background of each concentration in a serial dilution of a 1:1 mixture of the other two test articles. To evaluate antibody synergy in the combinations, the observed combination response matrix of pseudovirus neutralization was used as input for the online SynergyFinder platform (4), where quadruplicate data points were input separately. The highest single agent (HSA) reference model was applied, which quantifies synergy as the excess over the maximum response of a single drug in the combination. Synergy between antibodies in each combination is reported as an overall synergy score (the average of observed synergy across the dose combination matrix) as well as a peak HSA score (the highest synergy score calculated across the dose combination matrix). Synergy scores of less than -10, between -10 and 10, and greater than 10 indicate antagonistic, additive, and synergistic antibody combinations, respectively. While peak HSA scores report on synergy at the most optimal combination concentrations, the overall synergy score is less affected by outlier data points.

### Phagocytosis assay

Assay was performed with antibodies diluted to 100 ug/mL in PBS + 1% BSA. Antibodies were subjected to overnight incubation on tube rotator at 4°C in the presence of bead-biotinylated antigen mixture, followed by 3 washes. THP-1 cells were pelleted, resuspended in serum-free RPM and then added to wells containing bead-antigen-antibody mixture. The bead-antigen-antibody-cells mixture was incubated with cells in CO2 incubator for 18 hours. After that, cells were fixed and immunostained. Flow cytometry was peformed on Attune NXT and the resulted data were analyzed using FlowJo Software.

### Activation of classical complement pathway

ELISA-based method to evaluate the activation of the classical complement pathway was adapted from (*31, 45*). Endotoxin-free ELISA plates were coated with wither RBD or Trimer soluble proteins diluted in endotoxin-free PBS (HyClone) overnight. Plates were blocked with endotoxin-free 2% gelatin solution (Sigma) and incubated with anti-Spike antibodies of interest for 1 hour at +4°C. Plates were washed 3 times with endotoxin-free GVB buffer with Ca^2+^ and Mg^2+^ (GVB++, Complement Technology) and incubated with normal human serum (Complement Technology) diluted to 1.25% in GVB++ buffer for 1.5 hours at +37°C on an orbital shaker. Reaction was stopped by a wash with ice-cold PBS. Cells with deposited complement components were stained with anti-C4 antisera (Complement Technology) and a secondary anti-goat-HRP antibody (SouthernBiotech). Plates were incubated with HRP substrate and a stop solution according to manufacturer’s instructions (ThermoFisher). Optical density was measured on EnSpire Plate Reader (PerkinElmer)

### Receptor competition assay

ELISA plates were coated with either REF, UK or SA variant of RBD (SinoBio) overnight and washed with PBS (3x). Single antibody or a 3-Ab cocktail added at equimolar concentrations were added simultaneously with soluble human ACE2 protein (at a concentration of ∼ EC80 of its normal binding to RBD protein) and incubated at +37C on an orbital shaker for 1 hour. Plates were washed (3x) and subsequently probed for ACE2 binding with anti-ACE2 antibody.

### Authentic virus neutralization assay

Antibody combinations starting at 30 µg/ml per antibody were serially diluted in Dulbecco’s Phosphate Buffered Saline (DPBS)(Gibco™) using half-log dilutions. Dilutions were prepared in triplicate for each antibody and plated in triplicate. Each dilution was incubated at 37℃ and 5% CO2 for 1 hour with 10^3^ plaque forming units/ml (PFU/ml) of each SARS-CoV-2 variant [isolate USAWA1/2020, hCoV-19/USA/CA_CDC_5574/2020, BEI #NR-54011 (B.1.1.7) and hCoV-19/South Africa/KRISP-K005325/2020, BEI #NR-54009 (B.1.351)]. Each virus stock was passaged once from starting material in Vero E6 cells prior to use. Controls included Dulbecco’s Modified Eagle Medium (DMEM) (Gibco™) containing 2% fetal bovine serum (Gibco™) and antibiotic-antimycotic (Gibco™) only as a negative control and 1000 PFU/ml SARS-CoV-2 incubated with DPBS. Two hundred microliters of each dilution or control were added to confluent monolayers of NR-596 Vero E6 cells in duplicate and incubated for 1 hour at 37°C and 5% CO2. The plates were gently rocked every 15 minutes to prevent monolayer drying. The monolayers were then overlaid with a 1:1 solution of 2.5% Avicel® RC-591 microcrystalline cellulose and carboxymethylcellulose sodium (DuPont Nutrition & Biosciences, Wilmington, DE) and 2X Modified Eagle Medium (Temin’s modification, Gibco™) supplemented with 2X antibiotic-antimycotic (Gibco™), 2X GlutaMAX (Gibco™) and 10% fetal bovine serum (Gibco™). Plates were incubated at 37°C and 5% CO2 for 2 days. The monolayers were fixed with 10% neutral buffered formalin and stained with 0.2% aqueous Gentian Violet (RICCA Chemicals, Arlington, TX) in 10% neutral buffered formalin for 30 min, followed by rinsing and plaque counting. The half maximal inhibitory concentrations (IC50) were calculated using GraphPad Prism 8.

#### Focus reduction neutralization assay

Serial dilutions of antibodies were incubated with 10^2^ focus-forming units (FFU) of WA1/2020 D614G, BA.1, or BA.1.1. for 1 h at 37°C. Antibody-virus complexes were added to Vero-TMPRSS2 cell monolayers in 96-well plates and incubated at 37°C for 1 h. Subsequently, cells were overlaid with 1% (w/v) methylcellulose in MEM. Plates were harvested 30 h (WA1/2020 D614G) or 72 h (BA.1 and BA.1.1) later by removing overlays and fixed with 4% PFA in PBS for 20 min at room temperature. Plates were washed and sequentially incubated with an oligoclonal pool (SARS2-02, -08, -09, -10, -11, -13, -14, -17, -20, -26, -27, -28, -31, -38, -41, -42, -44, -49, -57, -62, -64, -65, -67, and -71 (*37*) of anti-S murine antibodies (including cross-reactive mAbs to SARS-CoV) and HRP-conjugated goat anti-mouse IgG (Sigma Cat # A8924) in PBS supplemented with 0.1% saponin and 0.1% bovine serum albumin. SARS-CoV-2-infected cell foci were visualized using TrueBlue peroxidase substrate (KPL) and quantitated on an ImmunoSpot microanalyzer (Cellular Technologies).

### Cryo-EM analysis of Trimer-Fab complexes

Cryo-EM analysis was performed at NovAliX (Strasbourg, France). Fabs were mixed with the SARS-CoV-2 S 6P trimer (6:1 molar ratio Fab per protomer) to a final Fab–S complex concentration of around 0.8 mg ml^−1^ and incubated at room temperature for 1H. Immediately before deposition of 3.5 μl of complex onto a 200 mesh, 1.2/1.3 C-Flat grid (protochips) that had been freshly glow-discharged for 30 sec at 3 mA using an ELMO (Cordouan). The sample was incubated on the grid for 15 s and then blotted with filter paper for 2 s in a temperature and humidity controlled Vitrobot Mark IV (T = 6 °C, humidity 100%, blot force 2) followed by vitrification in 100% liquid ethane. Single-particle cryo-EM data were collected on a Glacios transmission electron microscope (Thermo Fisher) operating at 200 kV. Movies were collected using EPU software for automated data collection. Data were collected at a nominal underfocus of −0.6 to −2.8 µm, at magnifications of 120,000× with a pixel size of 1.2 Å. Micrographs were recorded as movie stacks on a Falcon III direct electron detector (Thermo Fisher); each movie stack was fractionated into 13 frames, for a total exposure of 1.5 s corresponding to an electron dose of 50 e−/Å2. Drift and gain correction and dose weighting were performed using MotionCorr2. A dose-weighted average image of the whole stack was used to determine the contrast transfer function with the software Gctf. The following workflow was processed using RELION 4.0. Ab-initio cryo-EM reconstruction was low-pass filtered to 60 Å and used as an initial reference for 3D classification. The following subclasses depicting high resolution features were selected for refinement with various number of particles. IMM202190: 3 from 8 subclasses, 173,541 particles; IMM2084L 2 from 6 subclasses, 62,150 particles; IMM20253: 2 from 6 subclasses for Trimer, 86,974 particles, and 1 from 6 for monomers, 40,489 particles. Atomic models from PDB:7E8C, PDB:6XLU, PDB:6XM5 or PDB:7NOH for the Spike Trimer and PDB:6TCQ for the Fabs were used as starting point. Models were then rigid body fitted to the density in Chimera.

### Cryo-EM analysis of IMM20184/253 Fabs-Spike complex

Cryo-EM analysis was performed at nanoimaging Services (San Diego, USA). Electron microscopy was performed using an FEI Titan Krios (Hillsboro, Oregon) transmission electron microscope operated at 300kV and equipped with a Gatan BioQuantum 1967 imaging filter and Gatan K3 Summit direct detector. Vitreous ice grids were clipped into cartridges, transferred into a cassette and then into the Krios autoloader, all while maintaining the grids at cryogenic temperature (below -170C°). Automated data-collection was carried out using Leginon software(*46*), where high magnification movies were acquired by selecting targets at a lower magnification(*46*). Dose-weighted movie frame alignment was done using MotionCor2 (*47*) or Full-frame or Patch motion correction in cryoSPARC (*48*) to account for stage drift and beam-induced motion. The contrast transfer function was estimated for each micrograph using CTFfind4, gCTF, or Patch CTF in cryoSPARC (*49, 50*). Individual particles were selected using automated picking protocols and extracted into particle stacks in either Relion (*51, 52*) or cryoSPARC. The particles may then be submitted to reference-free 2D alignment and classification in either Relion or cryoSPARC. One dataset was collected for sample S Protein RBD + IMM20253 Fab + IMM20184 Fab, totaling 3,148 high magnification images. About 1.4M particles were selected from 1,666 manually curated micrographs using cryoSPARC 3.3 live. All subsequent data processing was carried out in cryoSPARC 3.3. These particles were subjected to three rounds of 2D classification, and about 300k good particles were selected. Second dataset with 30° tilt was collected for sample S Protein RBD + IMM20253 Fab + IMM20184 Fab, totaling 629 high magnification images. About 100k particles were selected from 244 manually curated micrographs using cryoSPARC 3.3 live. Third dataset with 30° tilt was collected for sample S Protein RBD + IMM20253 Fab + IMM20184 Fab, totaling 1,369 high magnification images. About 420k particles were selected from 1,055 manually curated micrographs using cryoSPARC 3.3 live. All subsequent data processing was carried out in cryoSPARC 3.3. These particles were subjected to one round of 2D classification. All micrographs from three sessions were combined for further processing in cryoSPARC 3.3. ∼1.9M particles were extracted and subjected to three rounds of 2D classification analysis and about 600k particles were selected for ab initio reconstruction. The particles were then subjected to three rounds of heterogenous refinement using the good and junk classes from ab initio reconstruction as reference volume. The good particles were selected and subjected to homogenous refinement and non-uniform refinement. The final 3D reconstruction was at a nominal resolution of 3.87 Å using ∼170K particles (see Sup. Figure 1E,F). The best 3D classes were submitted to homogeneous 3D refinement that includes dynamic masking. Reported resolutions were based on the gold standard FSC = 0.143 criterion. Maps were visualized using Chimera.

## ACKNOWLEDMENTS

This study was funded by the U.S. Department of Defense (DOD) Joint Program Executive Office for Chemical, Biological, Radiological and Nuclear Defense’s (JPEO-CBRND) Joint Project Manager for Chemical, Biological, Radiological and Nuclear Medical (JPEO-CBRN Medical), in collaboration with the Defense Health Agency (DHA), under contract W911QY2090019. The opinions, interpretations, conclusions and recommendations are those of the authors and are not necessarily endorsed by the U.S. Army.

The following reagents were obtained through BEI Resources, NIAID, NIH: VERO C1008 (E6), Kidney (African green monkey), Working Cell Bank, NR-596; SARS-CoV-2, Isolate hCoV-19/USA/CA_CDC_5574/2020, NR-54011 (deposited by Centers for Disease Control and Prevention) and SARS-CoV-2, Isolate hCoV-19/South Africa/KRISP-K005325/2020, NR-54009 (contributed by Alex Sigal and Tulio de Oliveira). The SARS-CoV-2 isolate USA-WA1/2020 starting material was provided by the World Reference Center for Emerging Viruses and Arboviruses (WRCEVA), with Natalie Thornburg as the CDC Principal Investigator.

## AUTHOR CONTRIBUTIONS

PAN, JMD, JPD, NBP, JLBS, BCH, NH, CN, AP, RJH, MN, HS, JPF, TS, NS, LGAM, ELS, RIJ, SMS, and LEM performed wet laboratory experiments. PAN, JMD, JPD, JLBS, AHN, AG, MSD, and MKR analyzed and interpreted the data. LFL, AG and DHG consulted and critically discussed the manuscript. PAN, JDM, PS, MJM and MKR supervised the project. PAN, JDM, JPD, JLBS and MKR wrote and AG, DHG, PS and MJM edited the manuscript. The manuscript was reviewed and cleared for publication by DoD’s JPEO-CBRN Medical, JPEO-CBRND.

## SUPPLEMENTARY MATERIALS

### Supplementary Tables

**Supplementary Table 1.** Breadth of binding of IMM20190/184/253 antibodies to RBD proteins bearing mutations found in CDC VOCs. EC50 (pM) relative to respective reference proteins measured using hTRF assay.

| Sort | His-tagged protein | Binding region | Variant | IMM20190 | IMM20253 | IMM20184 |
| --- | --- | --- | --- | --- | --- | --- |
| 1 | A352S | RBD |  | 25.4 | 33.0 | 27.7 |
| 2 | A475V | RBD | LY escape-3 | 70.2 | 34.6 | 30.3 |
| 3 | E406Q | RBD |  | 67.2 | 44.1 | 29.2 |
| 4 | E484K | RBD | South Africa (beta), Brazil (gamma) | 39.2 | 32.7 | 23.2 |
| 5 | E484Q | RBD | India (B.1.617) | 59.9 | 22.7 | 21.0 |
| 6 | F486S | RBD |  | 41.2 | 26.3 | 23.0 |
| 7 | F490S | RBD | Peru (lambda) | 14.7 | 73.4 | 60.8 |
| 8 | K417N | RBD | South Africa (beta), India (delta+) | >500 | 67.4 | 44.3 |
| 9 | K417N, E484K, N501Y | RBD | South Africa (beta) | not detected | 25.0 | 20.7 |
| 10 | K417T, E484K, N501Y | RBD | Brazil (gamma) | >500 | 22.6 | 17.9 |
| 11 | K444R | RBD |  | 33.9 | 26.7 | 20.9 |
| 12 | L452R | RBD | USA (epsilon), India (all) | 40.9 | 34.3 | 41.2 |
| 13 | L452R, T478K | RBD | India (B.1.617.2, delta) | 52.8 | 61.7 | 111 |
| 14 | N439K | RBD | Scotland, Europe | 76.6 | 56.5 | 52.1 |
| 15 | N440K | RBD |  | 45.6 | 39.7 | 32.1 |
| 16 | N501Y | RBD | UK (alpha), SA (beta), Brazil gamma) | 311 | 59.0 | 42.2 |
| 17 | Spike RBD (319-591) | RBD | Wuhan / Washington reference | 69.9 | 53.4 | 46.2 |
| 18 | T478I | RBD |  | 16.4 | 45.5 | 68.5 |
| 19 | Y453F | RBD | Denmark (mink) | 71.6 | 65.6 | 50.1 |
| 20 | A222S, D614G | S1 | Europe, Spain | 51.0 | 45.1 | 29.9 |
| 21 | D614G | S1 | Multiple | 69.9 | 62.9 | 44.1 |
| 22 | K417N, E484K, N501Y, D614G | S1 | South Africa (beta) | 391 | 34.7 | 25.6 |
| 23 | Spike S1 (WT) | S1 | Wuhan / Washington reference | 58.8 | 50.0 | 34.4 |
| 24 | SARS-CoV-1 | S1 | Wild type | not detected | 137.0 | >500 |
| 25 | T19R, G142D, E156G, Δ157-158, L452R, T478K, D614G, P681R | S1 | India (B.1.617.2, delta) | 22.2 | 40.9 | 30.6 |
| 26 | E154K, L452R, E484Q, D614G, P681R | S1 | India (B.1.617.1, kappa) | 38.2 | 49.2 | 22.7 |
| 27 | ΔHV69/70, Y453F, D614G | S1 | Denmark (mink) | 48.1 | 67.1 | 34.8 |
| 28 | ΔHV69/70, ΔY144, N501Y, A570D, D614G, P681H | S1 | UK (alpha) | >500 | 63.1 | 49.1 |

### Supplementary Figures

**Supplementary Figure 1.**
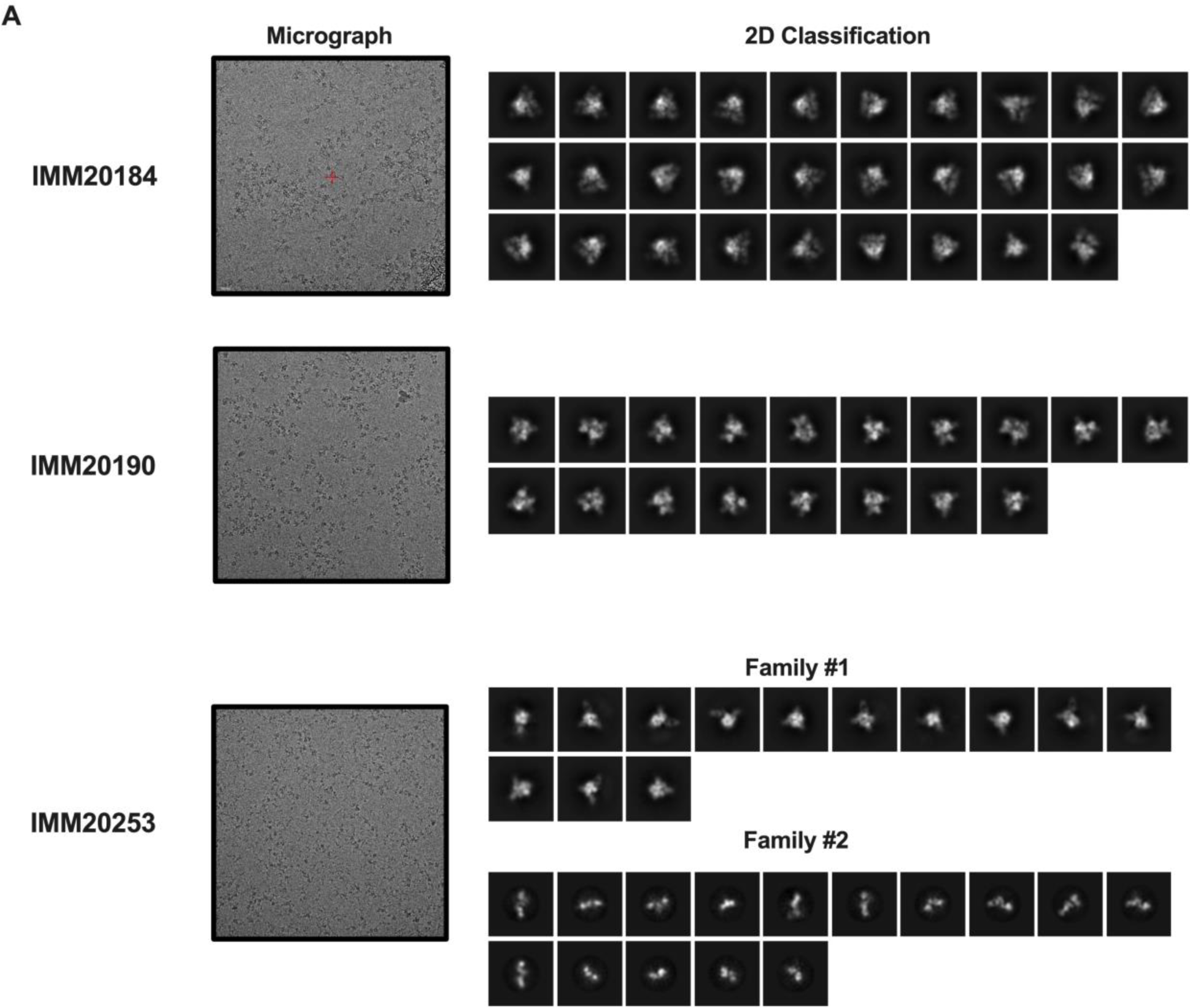
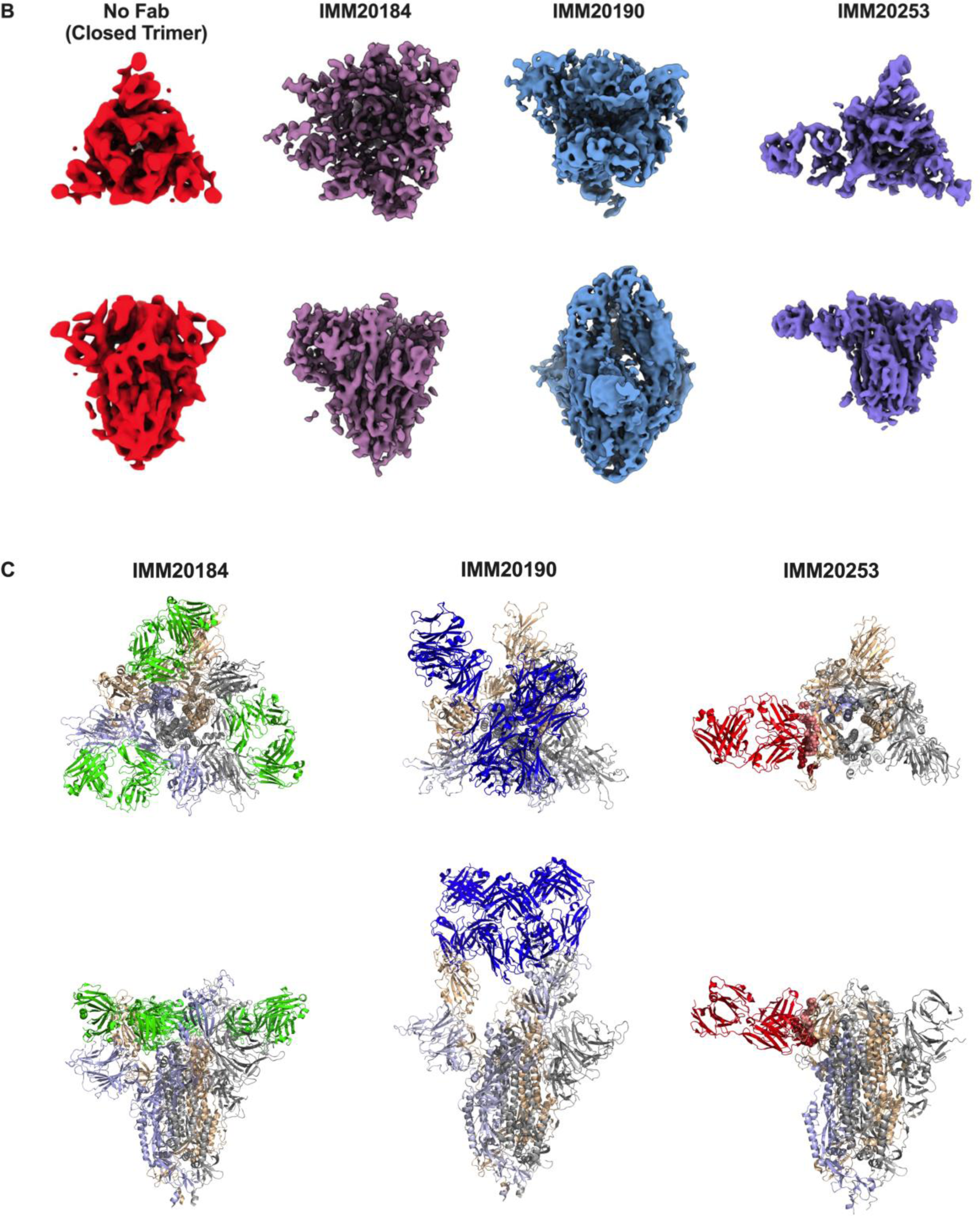
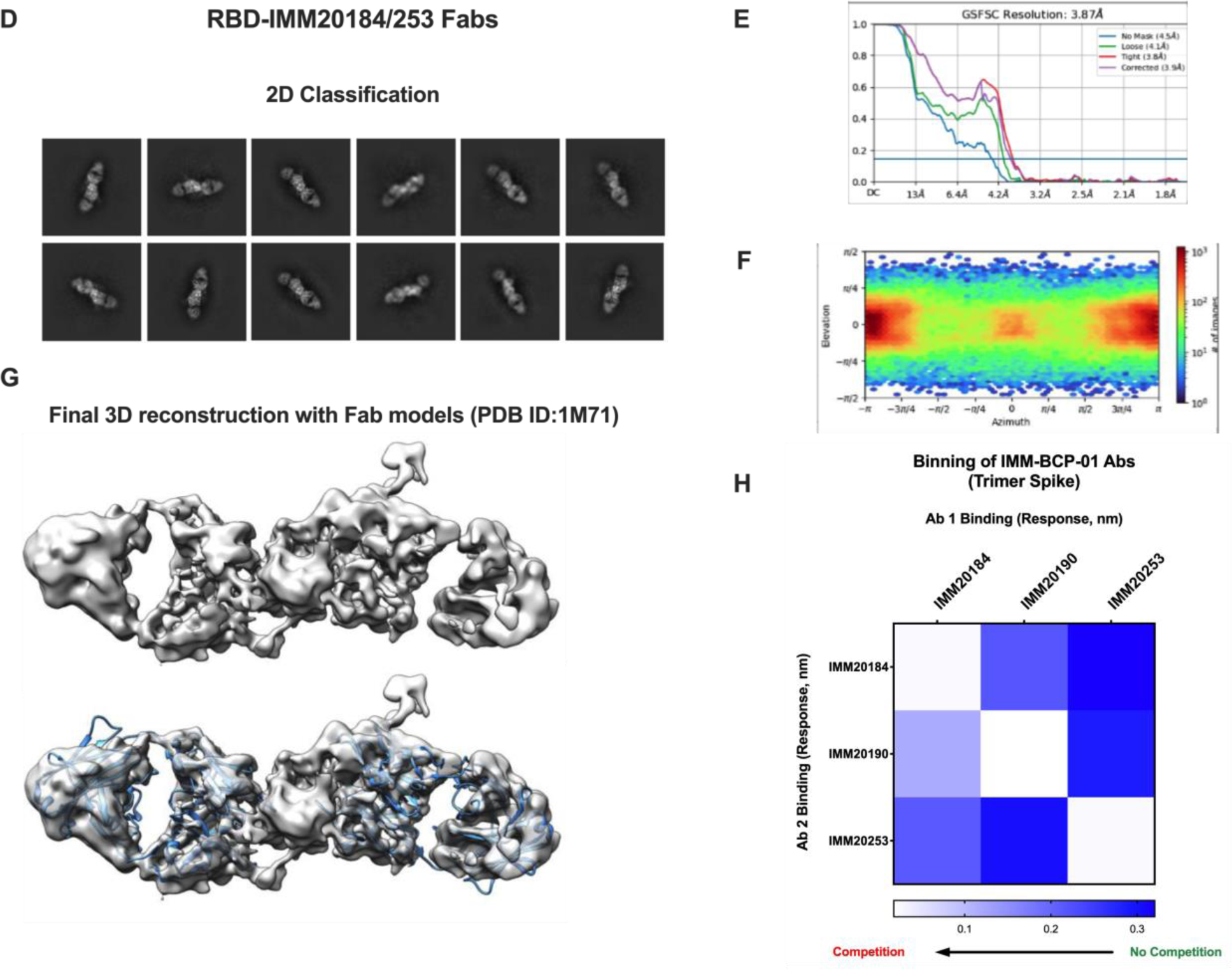
Cryo-EM micrographs reveal sites of IMM20184/20190/20253 Fabs binding to Spike protein. (A) Cryo-EM micrographs and 2D classification of Trimer-Fab complexes shown in Figure 1A. IMM20253 binding to Trimer generates two families (shown as Family#1 and 2). (B). Comparison of 3D reconstruction data (density only) for a closed Trimer conformation, IMM20184 Fab-Trimer, IMM20190 Fab-Trimer and IMM20253 Fab-Trimer complexes in support of data shown in Figure 1A. (C) Models PDB:7E8C, PDB:6XLU, PDB:6XM5 or PDB:7NOH for Trimer and PDB:6TCQ for the Fabs demonstrate binding patterns and attack angles for IMM20184/190/253 antibodies. (D) CryoEM micrographs and 2D classification of a IMM20184 Fab – RBD – IMM20253 Fab complex. Simultaneous binding of both Fabs is clearly visible. (E) Fourier shell correlation (FSC) curves of the final 3D refinement of data from panel D in cryoSPARC 3.3 for different types of masks. The resolution of the final map was calculated based on a FSC of 0.143. (F). Viewing directional distribution for the final refinement run for the complex shown in panels D and E, generated by cryoSPARC 3.3. The viewing direction distribution histogram shows the number of images with a particular viewing direction at each (elevation, azimuth angle). (G). Final 3D reconstruction of data shown in Supp. Figure 1D and Figure 1B. Primary data (density) and modelling using Fab model PDB:1M71 are shown. (H). Antibody binning on Octet Qke. IMM20184, IMM20190 and IMM20253 do not compete for soluble Trimer and RBD protein binding. Heat map values represent binding of the first antibody to Trimer (top), followed by binding of the second antibody (left), measured as Response parameter in nm.

**Supplementary Figure 2.**
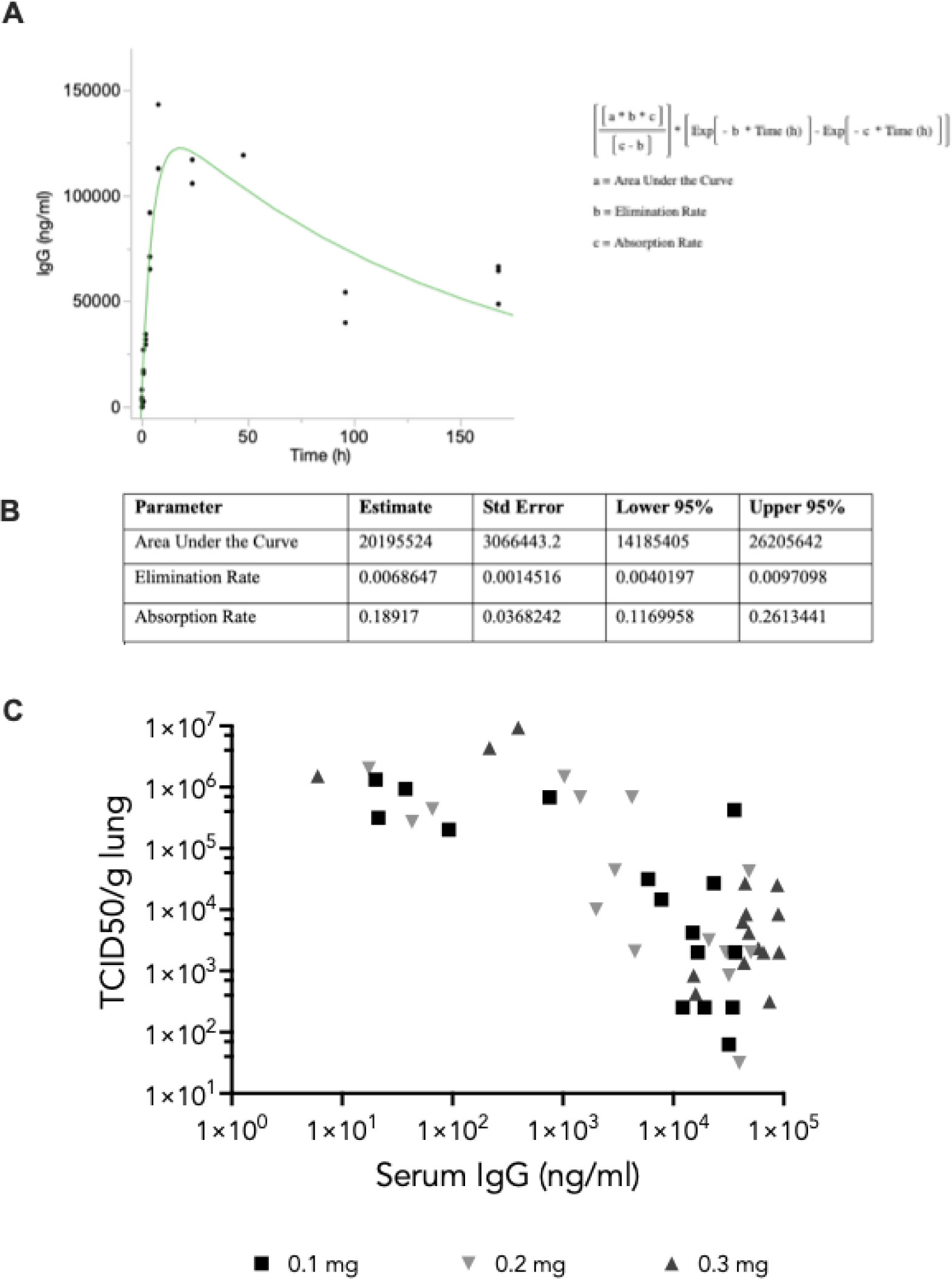
Antibody exposure and pharmacokinetics in dosed hamsters. (A) Pharmacokinetics of the 3-Ab cocktail in hamsters. The 3-Ab cocktail (0.3 mg each) was administered i.p. into Syrian Golden hamsters and terminal bleeds (n = 4 per time point) were taken at 0.25, 0.5. 1, 2, 4, 8, 24, 48, 96, and 168 hours post administration. Total human IgG levels were determined by anti-human ELISA. Pharmacokinetics in animals exhibiting < 1000 ng/mL IgG in serum at timepoints >30 minutes post-injection. Green line is the calculated curve using the formula shown on the right. (B) PK parameters of data from panel A. (C) Viral titer in lungs of infected hamsters depends upon Ab exposure. Syrian golden hamsters challenged with 3.3x10^5^ TCID_50_ viral inoculation of a non-adapted WA_CDC-WA1/2020 SARS-CoV-2 isolate were treated with 3-Ab cocktail (IMM20184/IMM20190/IMM20253), at various dose levels, six hours post inoculation with virus. Lungs were harvested at day 4 post-treatment and viral titers were determined by TCID_50_ assay. Terminal levels of IgG in blood were quantified by anti-human ELISA.

**Supplementary Figure 3.**
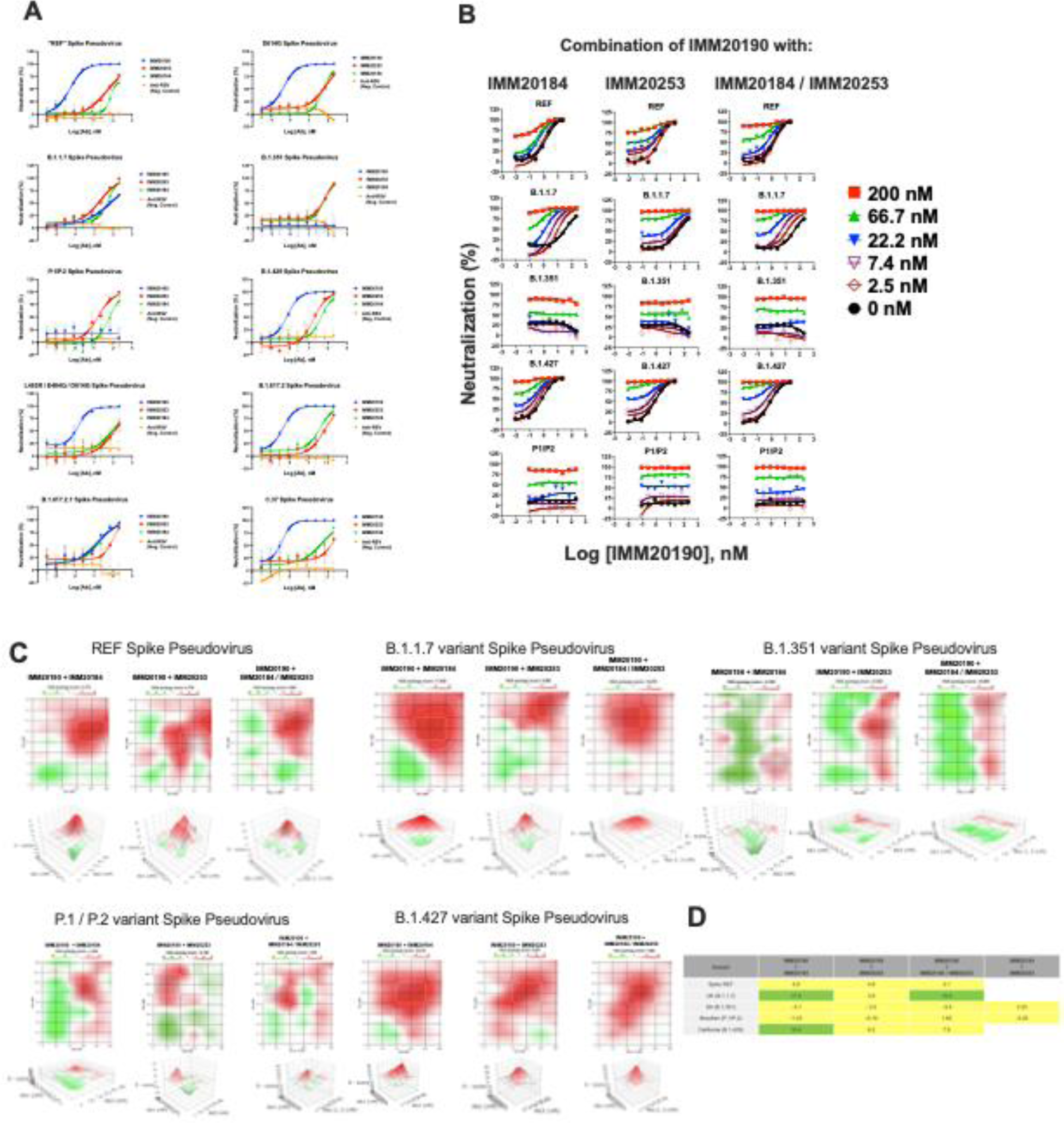
Three selected antibodies have a synergistic neutralizing effect. (A) Neutralization properties of standalone IMM20190, IMM20184 and IMM20253 antibodies against 10 different Spike variant pseudoviruses. (B) REF, B.1.1.7 (alpha), B.1.351 (beta), P1 (gamma), and B.1.427 (epsilon) pseudovirus variant neutralization by IMM20190 combination with either IMM20184, IMM20253 or both. (C) The Highest Single Agent (HSA) scores for 2-Ab and 3-Ab combinations. IMM20190 was mixed with IMM20184 and IMM20253 at 1:0.5:0.5 ratio. (D) HSA scores for two and three antibody cocktail.

**Supplementary Figure 4.**
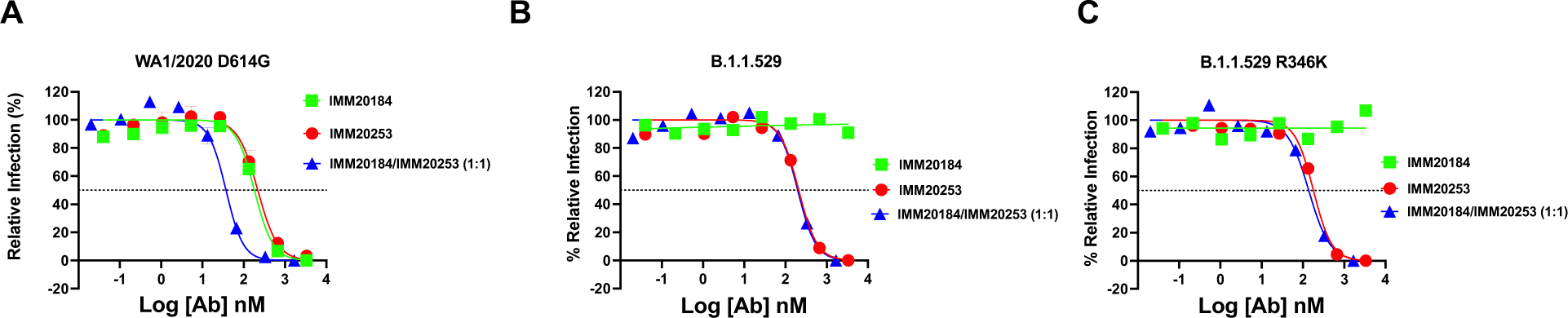
Focus reduction neutralization assay (FRNT) of SARS-CoV-2 variants in the presence of IMM20184, IMM20253 and IMM20253/184 combination. (A) Relative infection of WA1/2020 D614G, (B) Omicron (BA.1) and (C) Omicron BA.1.1 virus variants in the presence of IMM20184, IMM20253 and IMM20184/253 antibodies. Data are representative of three independent experiments performed in duplicate.

**Supplementary Figure 5.**
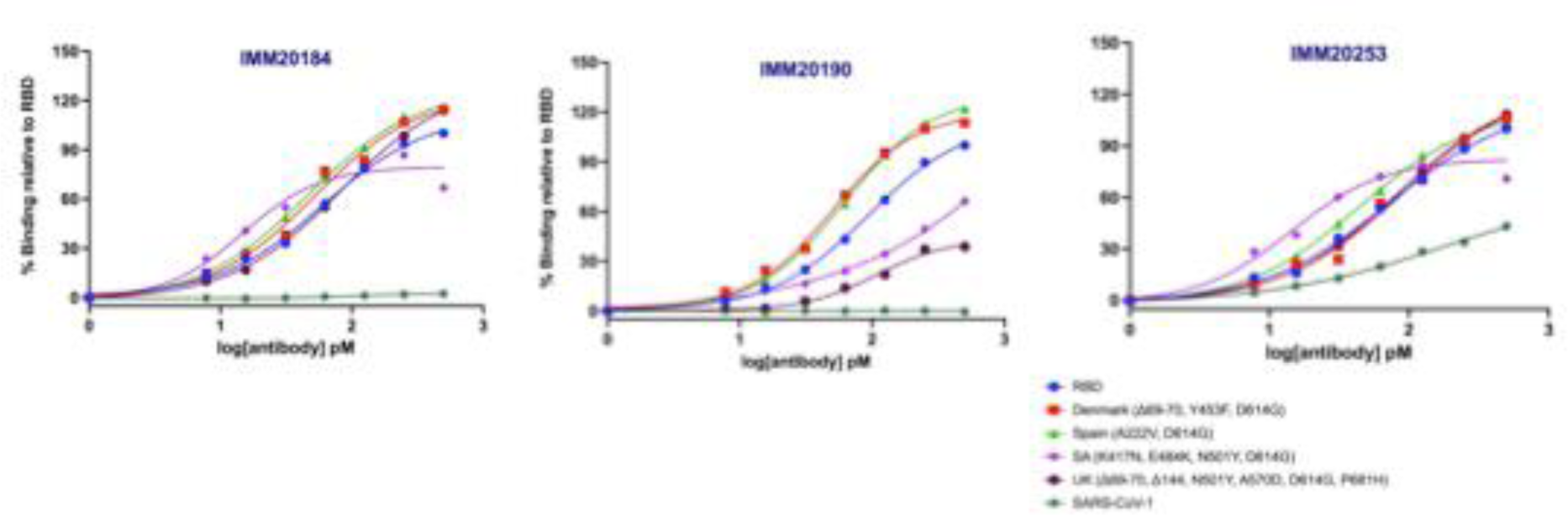
Binding of IMM antibodies to soluble RBD proteins from SARS-CoV-1 and SARS-CoV-2 variants in a steady-state hTRF assay.

